# Host-dependent induction of disease tolerance to infection by tetracycline antibiotics

**DOI:** 10.1101/833269

**Authors:** Henrique G. Colaço, André Barros, Ana Neves-Costa, Elsa Seixas, Dora Pedroso, Tiago R. Velho, Katharina Willmann, Hyon-Seung Yi, Minho Shong, Vladimir Benes, Sebastian Weis, Thomas Köcher, Luís F. Moita

**Affiliations:** Innate Immunity and Inflammation Laboratory, Instituto Gulbenkian de Ciência, Rua da Quinta Grande 6, 2780-156 Oeiras, Portugal; Research Center for Endocrine and Metabolic Diseases, Chungnam National University School of Medicine, Daejeon 35015, Korea; EMBL Genomics Core Facilities, D-69117 Heidelberg, Germany; Institute for Infectious Disease and Infection Control, Jena University Hospital; Department of Anesthesiology and Intensive Care Medicine, Jena University Hospital; Center for Sepsis Control and Care, Jena University Hospital, 07747 Jena, Germany; Vienna BioCenter Core Facilities GmbH, 1030 Vienna, Austria; Instituto de Histologia e Biologia do Desenvolvimento, Faculdade de Medicina, Universidade de Lisboa, Portugal

**Author notes:** Corresponding author: Luis F. Moita.

## Abstract

Synergy of resistance and disease tolerance mechanisms is necessary for an effective immune response leading to survival and return to homeostasis when an organism is challenged by infection. Antibiotics are used for their resistance enhancement capabilities by decreasing pathogen load, but several classes have long been known to have beneficial effects that cannot be explained strictly on the basis of their capacity to control the infectious agent. Here we report that tetracycline antibiotics, a class of ribosome-targeting drugs, robustly protects against sepsis by inducing disease tolerance, independently from their direct antibiotic properties. Mechanistically, we find that mitochondrial inhibition of protein synthesis perturbs the electron transfer chain and leads to improved damage repair in the lung and fatty acid oxidation and glucocorticoid sensitivity in the liver. Using a partial and acute deletion of *CRIF1* in the liver, a critical mitoribosomal component for protein synthesis, we find that mice are protected against bacterial sepsis, an observation which is phenocopied by the transient inhibition of complex I of ETC by phenformin. Together, we demonstrate that ribosome-targeting antibiotics are beneficial beyond their antibacterial activity and that mitochondrial protein synthesis inhibition leading to ETC perturbation is a novel mechanism for the induction of disease tolerance.

## Introduction

Organismal survival and optimal function require robust homeostatic responses to variable and challenging environmental conditions (Rajan and Perrimon, 2011). Understanding core physiological principles in response to infection and its genetic circuitry are current central questions in biology (Chovatiya and Medzhitov, 2014). Sepsis is a prime example of extreme homeostasis disruption caused by infection and therefore constitutes an excellent model to study homeostatic circuits and inter-organ communication principles (Colaço and Moita, 2016). Sepsis is a complex disorder caused by a non-adaptive host response to an infection (Singer et al., 2016), leading to acute organ dysfunction and consequent high risk of death (Cecconi et al., 2018). It is the leading cause of death in intensive care units and the third cause of overall hospital mortality (Van Vught et al., 2016). The pathophysiology and molecular bases of sepsis remain poorly understood. The urgently needed novel therapies for sepsis can only be inspired by new insights into the molecular bases of multi-organ failure and endogenous tissue protective mechanisms. Hallmarks of sepsis include an acute burst in pro-inflammatory cytokine production (Wiersinga et al., 2014) and metabolic failure (Wyngene et al.), both leading to severe tissue damage and high mortality rates. Current management of sepsis patients is limited to control of infection with antibiotics and organ support measures, with most attempts to modulate immune response resulting in failure (Cohen et al., 2015).

Surviving a severe infection requires the synergy between two evolutionarily conserved defense strategies that can limit host disease severity. Resistance relies on extensively explored pathogen recognition, signalling transduction pathways, and effector mechanisms to reduce pathogen load, while disease tolerance provides host tissue damage control and limits disease severity irrespectively of pathogen load (Soares et al., 2017) using mechanisms that have only recently begun to be explored at the molecular, cellular and organismal levels (Soares et al., 2017). Resistance mechanisms – which leads to pathogen detection, containment and elimination – are not enough to guarantee recovery from infection. In fact, many sepsis patients die (currently ∼25%) despite effective eradication of the inciting pathogen. Therefore, disease tolerance based therapies are likely to be critical adjuvant components of sepsis treatment (Figueiredo et al., 2013; Ganeshan et al., 2019; Weis et al., 2017).

All eukaryotic organisms are equipped with surveillance mechanisms to detect and correct perturbations in homeostasis. Organelle dysfunction caused by pathogens, toxins, drugs, physical insults or nutritional changes are rapidly communicated to the nucleus where a transcriptional response initiates a compensatory response (Sawa et al., 2016). Such stress responses are critical for the initiation of an effective immune response (Colaço and Moita, 2016) and cytoprotective-based lifespan extension programs (Shore and Ruvkun, 2013). Remarkably, locally induced cytoprotective stress responses can be communicated to distant organs, generating systemic beneficial effects (Durieux et al., 2011; Owusu-Ansah et al., 2013).

Mitochondria have essential roles in bioenergetics, metabolism, and cell signaling. They have evolved complex but still poorly understood surveillance programs to monitor physiological perturbations leading to the initiation of cytoprotection mechanisms (Yun and Finkel, 2014). Pioneer work in *C. elegans* revealed that mild perturbations in mitochondrial function induced both by genetic defects in the electron transfer chain (ETC) (Dillin et al., 2002) or inhibition of mitochondrial protein synthesis (Houtkooper et al., 2013) resulted in extended lifespan. In mice, several studies have documented metabolic benefits arising from inhibition of ETC activity in the context of obesity and insulin resistance (Chung et al., 2017; Masand et al., 2018; Pospisilik et al., 2007) with no significant effects in longevity (Deepa et al., 2018). While the molecular mechanisms of mitochondrial stress responses remain poorly understood in mammals, it is generally accepted that perturbations in mitochondrial function involve: 1) retrograde signaling to the nucleus, which activates a transcriptional program known as the mitochondrial unfolded protein response (UPR^mt^) (Fiorese et al., 2016; Houtkooper et al., 2013), and 2) metabolic adaptation, which may increase fitness in adverse conditions (Masand et al., 2018). Besides mitochondria, perturbation of other core cellular functions such as insulin/insulin growth-factor signaling (Kenyon et al., 1993), mRNA translation (Govindan et al., 2015), and ER protein homeostasis (Fouillet et al., 2012) have shown beneficial effects in numerous experimental models.

We have recently shown that activation of DNA damage responses in response to low level of DNA damage caused by anthracyclines induces disease tolerance and promotes survival in several sepsis mouse models (Figueiredo et al., 2013). In this study, we set out to identify novel protective mechanisms against infection activated by mild disruption of core cellular functions. We found that protein synthesis inhibition by ribosome-targeting antibiotics like doxycycline, robustly and reproducibly increase survival independently of its antibiotic effects. These results are associated with changes in mitochondrial function, fatty acid metabolism and response to glucocorticoids, which can be phenocopied by mild and transient chemical and genetically induced perturbations of mitochondrial ETC.

## Results

### Doxycycline confers protection in a mouse model of bacterial sepsis

To identify novel disease tolerance mechanisms initiated to resolve perturbation of core cellular functions, in addition to DNA damage, we selected a panel of clinically approved drugs known to interfere with basic cellular functions and pathways. These drugs were intraperitoneally injected into C57BL/6J mice at time of infection with an *E. coli* strain carrying resistance to tetracyclines (TetR) and chloramphenicol (CamR) to control for the effects of ribosomal-targeting antibiotics directly on bacterial viability. From the panel of tested drugs, treatment with low-dose doxycycline, a tetracycline antibiotic known to bind to the mitochondrial ribosome and block translation of mitochondrial-encoded mRNA (Lin et al., 2018), revealed a robust and reproducible increase in survival in multiple independent experiments (Figure 1A). We used rectal temperature and body weight measurements to assess disease severity and found that doxycycline-treated mice had less severe hypothermia than PBS-treated controls and increased body temperature close to normal levels within 48h of infection (Figure 1B). Marginal but reproducible and consistent differences in body weight were found between groups, with doxycycline-treated mice presenting slightly lower body weight at 48h (Figure 1C), a difference not found in the absence of infection (data not shown). Notably, treatment with chloramphenicol, another but structurally unrelated ribosome-targeting antibiotic working by mitochondrial mRNA translation inhibition, also confers protection in this sepsis model (Figure S1A, 1B, 1C).

**Figure 1.**
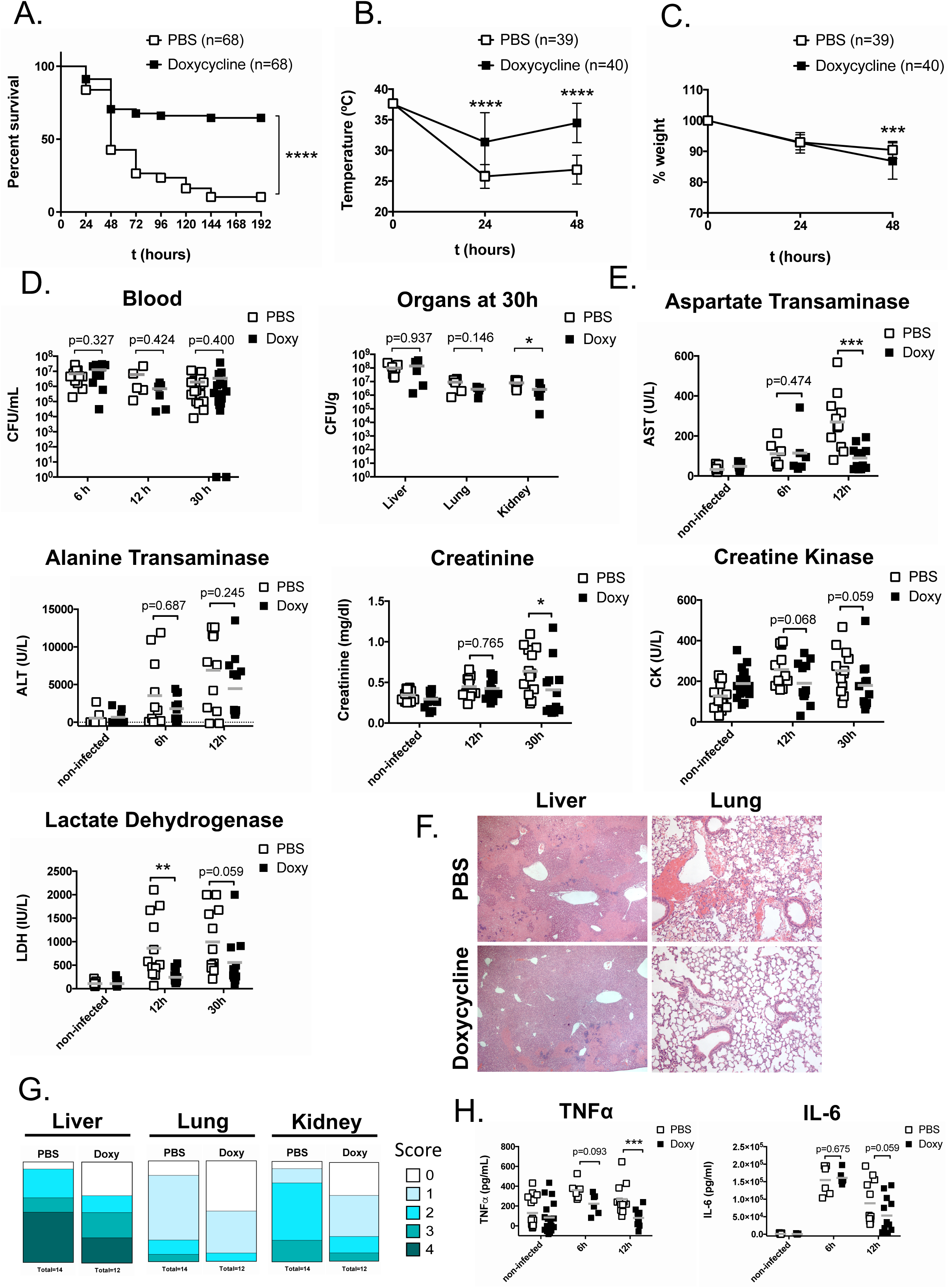
Doxycycline confers protection in a mouse model of bacterial sepsis. Survival (A), rectal temperature (B) and % initial body weight (C) after infection of C57BL/6J mice with 3×10^8^ CFU/mouse TetR CamR *E. coli* and treatment with 1.75 μg/g body weight doxycycline (or PBS as a control) at 0, 24 and 48 h. (D) Bacterial load in mouse blood, liver, lung, and kidney at the indicated time-points after infection. (E) Levels of the organ damage markers Aspartate Transaminase (AST), Alanine Transaminase (ALT), Creatinine, Creatine Kinase (CK), and Lactate Dehydrogenase (LDH) at the indicated time-points after infection. (F) Representative images of Hematoxylin-Eosin stained liver and lung 30 h after infection. (G) Organ damage score in Hematoxylin-Eosin stained slides from liver, lung, and kidney 30 h after infection. Score 0 = no lesions; 1 = very mild; 2 = mild; 3 = moderate; 4 = severe lesions. (H) TNFα and IL-6 levels in mouse serum at the indicated time-points after infection. (A) represents pooled data from twelve independent experiments. (B) and (C) represent mean±SD pooled from six independent experiments. (D), (E), and (H) represent pooled data from at least two independent experiments; squares represent individual mice and gray bars indicate the mean. See also Figure S1.

We found no differences in the bacterial loads of doxycycline-treated mice in either blood (Figure 1D, left panel) or tissue samples (Figure 1D, right panel) collected at 6, 12, and 30h after infection, except for a modest reduction in kidney CFUs only at 30h (Figure 1D, right panel). These results suggest that doxycycline induces disease tolerance by limiting disease severity without affecting pathogen load (Soares et al., 2017), independently of a direct antibiotic effect and modulation of host resistance mechanisms acting on the TetR *E. coli* strain to clear the infection. Remarkably, despite similar bacterial burden between groups, doxycycline treated mice show reduced levels of tissue damage in the major target organs of sepsis (Figure 1E, 1F, 1G). Serum levels of the liver damage markers aspartate transaminase (AST) and alanine transaminase (ALT) determined at different time-points after infection revealed very significantly reduced AST levels at 12h, with no significant differences in ALT levels (Figure 1E). The kidney damage marker creatinine was markedly reduced at 30h, whereas the muscle damage marker creatine kinase (CK) showed modest but non-statistically significant differences (Figure 1E). Lactate dehydrogenase, an unspecific damage marker, showed markedly reduced levels at 12h (Figure 1E). Reduced tissue damage was also documented in a blind histopathology analysis of liver, lung, and kidney, in which tissues were scored for the presence and dimension of necrotic areas as well as leukocyte infiltration (Figure 1F, 1G). In all analyzed tissues, doxycycline-treated mice globally scored lower for damage, including a higher number of animals with no visible damage (score 0) (Figure 1G). These changes were more pronounced in the liver, where necrotic areas of doxycycline-treated mice were markedly reduced, and in the lung, where we observed reduced neutrophil infiltration, hemorrhage, and thickening of the alveolar wall upon doxycycline treatment (Figure 1F). We then analyzed the levels of the pro-inflammatory cytokines tumor necrosis factor α (TNFα) and interleukin 6 (IL-6), which are quickly and strongly increased during sepsis. In serum of *E. coli*-infected mice, TNFα levels were found significantly reduced at 12h, while IL-6 levels showed slight, non-statistically significant reduction (Figure 1H). However, in mouse bone marrow-derived macrophages (BMDM) treated with doxycycline and stimulated with *E. coli*, no differences were found in cytokine levels (Figure S1D), suggesting that doxycycline does not directly modulate the inflammatory response during sepsis, including at the transcriptional level, but point instead to the role of the drug in tissue protection mechanisms that induce disease tolerance.

To test the specificity of doxycycline-induced disease tolerance across infection models, we used a model of systemic fungal infection induced by intravenous injection of *Candida albicans* and found no differences in survival rates of treated mice (Figure S1E). Similarly, in a model of cerebral malaria induced by *Plamodium berghei* Anka no differences were found in survival rates (Figure S1F) despite reduced percentage of infected red blood cells upon doxycycline treatment (Figure S1G), in line with the well-established anti-malarial effects of the drug (Gaillard et al., 2015). These results suggest that doxycycline-induced mechanisms of disease tolerance are specific for bacterial infections.

### Doxycycline improves lung pathology by inducing a tissue repair response

Acute lung injury is a common feature of sepsis and a major contributor for mortality and morbidity of the patients (Sadowitz et al., 2011). Guided by the improvement of lung pathology upon doxycycline treatment (Figure 1F, 1G), we explored the effect of local administration of the drug to the lung. We found that a single intra-tracheal administration of doxycycline 2 hours before *E. coli* infection results in significant increase in survival (Figure 2A), supporting a central role of the lung in sepsis outcome, in agreement with our previous findings in the case of anthracyclines (Figueiredo et al., 2013). To further explore the protective role of doxycycline in the lung, we used a model of Influenza infection which causes lung damage and inflammation. To that end, mice were intranasally challenged with a sublethal dose (100 pfu/mouse) of Influenza PR/8 and doxycycline was injected intraperitoneally at days 4, 5, and 6 post-infection, the period when viral loads are higher and lung lesions become apparent. We observed that doxycycline-treated mice lose body weight at a similar pace to the controls, but recover faster from day 5 post-infection onwards, with a significant difference in body weight at day 10 post-infection (Figure 2B). These results suggest that doxycycline may trigger tissue repair mechanisms at the level of the lung, which result in faster recovery.

**Figure 2.**
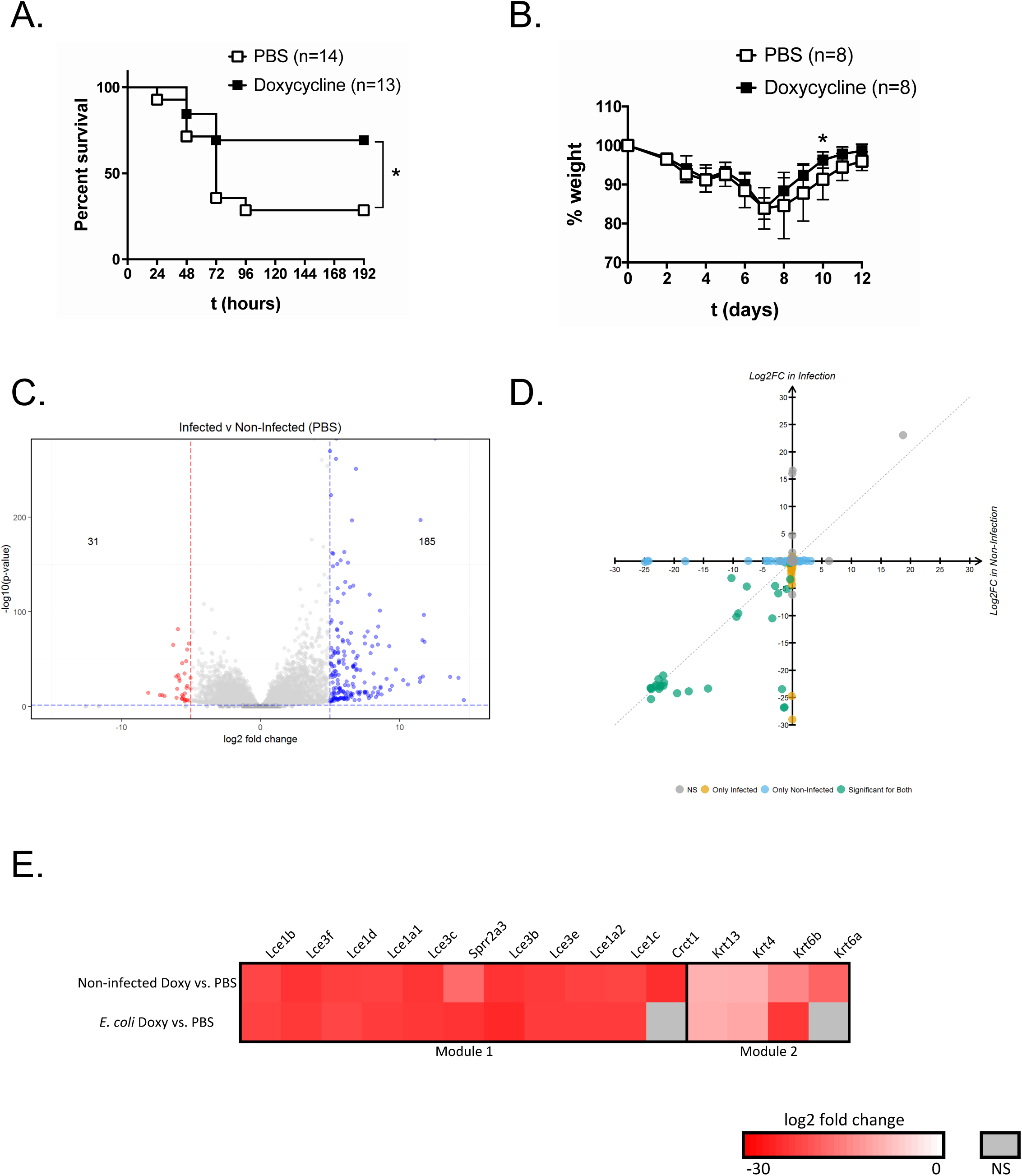
Doxycycline improves lung pathology by reprogramming basal cells. (A) Survival of mice after intra-tracheal delivery of 1.75 μg/g body weight doxycycline (or PBS as a control) followed by infection with *E. coli* 2h later. (B) Percentage of initial weight in mice infected with a sublethal (100 pfu/mouse) dose of Influenza A PR/8 and treated with 1.75 μg/g body weight doxycycline (or PBS as a control) at days 4, 5, and 6 post-infection. (C) Volcano plot with differential expression of genes in *E. coli*-infected and non-infected, PBS-treated mice from bulk RNA-seq analysis in the lung at 12 h. Numbers indicate genes with log2 fold change <-5 or >5 and p<0.05. (D) Scatter plot of genes affected by doxycycline treatment in infected versus non-infected groups. Yellow dots indicate genes differentially expressed in infected mice (p<0.05); blue dots indicate genes differentially expressed in non-infected mice (p<0.05); green dots indicate genes differentially expressed in both conditions (p<0.05); gray dots indicate non-statistically significant genes (p≥0.05). (E) Heat maps of genes affected by doxycycline treatment in non-infected mice (log_2_ fold change <-5; p<0.05), after clustering with DAVID ‘Gene Functional Classification’. (A) represents pooled data from two independent experiments. (B) is representative of two independent experiments. See also Figure S2.

To investigate possible mechanisms leading to epithelial repair, we performed bulk RNA-seq in mouse lungs 12h after infection with *E. coli* and treatment with doxycycline and compared with non-infected and PBS-treated controls. We found a large number of expected genes up-regulated upon infection (Figure 2C), mostly related to an acute inflammatory response (Figure S2A). Doxycycline treatment did not change the expression of the majority of these genes, but instead resulted in the strongly reduced expression of a small group of genes in both infected and non-infected groups (Figure 2D, Figure S2B), suggesting that drug-induced changes are independent of the infection. Functional clustering analysis in non-infected, doxycycline-treated mice showed a remarkable similarity in the function of down-regulated genes, with 60% of the genes clustering in pathways related to keratinization and epithelium differentiation (Figure 2E, Figure S2C). In particular, the basal cell markers Krt6a and Krt6b are severely down-regulated (Figure 2E), suggesting that doxycycline might be driving differentiation of lung progenitor cells (Hackett et al., 2011), leading to a more effective repair of the lung epithelium. To more directly test this possibility, we have used diphteria toxin to deplete basal cells from mice expressing the diphteria toxin receptor under a Krt6a promoter (Krt6a-DTR mice) (Zuo et al., 2015). Krt6a-DTR mice infected with *E. coli* as previously described can still be substantially protected by doxycycline treatment (Figure S2D), suggesting that basal cells are the unlikely single target of the drug and that the enhanced lung repair capacity induced by doxycycline depends on the combination of its effect on multiple cell lineages, rather than basal cells alone.

### Doxycycline improves liver pathology

We then turned our attention to the liver as doxycycline strongly limits tissue damage in this organ in response to infection (Figure 1D-G). To begin exploring the effect of doxycycline on liver function in sepsis, we performed bulk RNA-Seq in mouse liver 8 hours after *E. coli* infection with and without doxycycline treatment. Similar to the lung, infection resulted in up-regulation of a high number of genes associated with an inflammatory response in the liver (Figure 3A and S3A). Surprisingly, when comparing PBS and doxycycline treated mice in the absence of infection, we found no statistically significant differential gene expression (Figure 3B), suggesting that the transcriptional signature of tissue repair found in the lung (Figure 2D and 2E) is not present in the liver. Functional analysis of *E. coli*-infected, doxycycline-treated mice revealed a discrete number of up- and down-regulated genes compared to PBS-treated controls (Figure 3B), with a single significant cluster that relates to decreased production of collagen in infected, doxycycline treated mice (Figure S3B). Collagen is a well-known marker of liver fibrosis and changes in collagen metabolism have been associated with disease severity in sepsis (Gäddnäs et al., 2009). These findings, together with serology and pathology data (Figure 1E, F and G) further support a role for doxycycline in limiting liver damage from very early time-points of infection.

**Figure 3.**
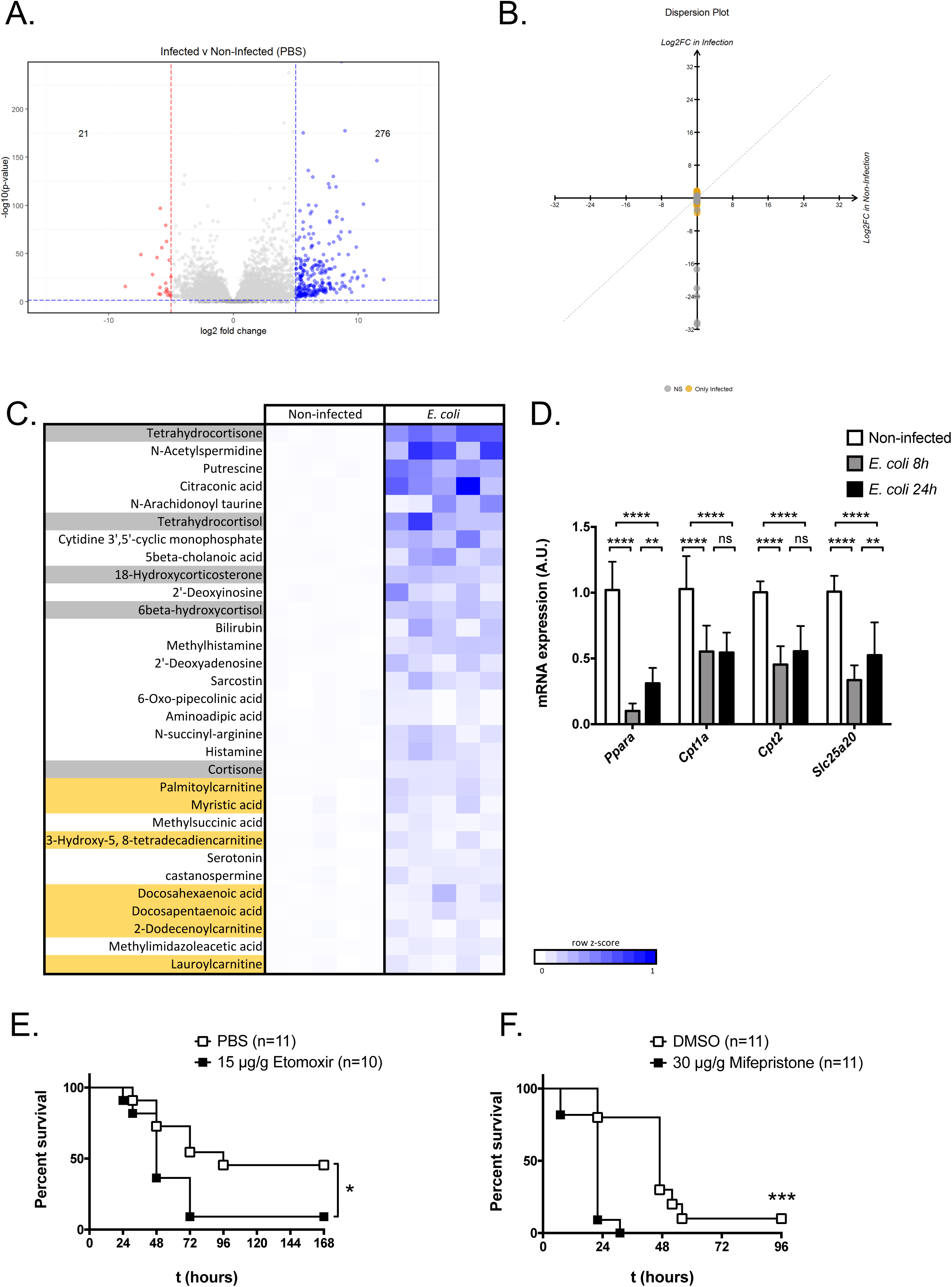
Fatty acid oxidation and response to glucocorticoids are essential for sepsis outcomes. (A) Volcano plot with differential expression of genes in infected and non-infected, PBS-treated mice from bulk RNA-seq analysis in the liver at 8 h. Numbers indicate genes with log2 fold change <-5 or >5 and p<0.05. (B) Scatter plot of genes affected by doxycycline treatment in infected versus non-infected groups. Yellow dots indicate genes differentially expressed in infected mice (p<0.05); gray dots indicate non-statistically significant genes (p≥0.05). (C) Untargeted metabolomics analysis in mouse liver 8h after infection. Top up-regulated metabolites in liver of infected mice (p<0.05). Glucocorticoids are highlighted in gray and metabolites involved in fatty acid oxidation are highlighted in yellow. (D) Expression of *Ppara* and several of its targets by qPCR in mouse liver at 8h and 24h post-infection. Data represent mean±SD of 5 mice assayed in triplicate. (E, F) Survival after infection of C57BL/6J mice with 3×10^8^ CFU/mouse TetR CamR *E. coli* and treatment with 15 μg/g body weight Etomoxir or PBS as a control (E) and 30 μg/g body weight Mifepristone or 100% DMSO as a control (F). Both graphs represent pooled data from two independent experiments. See also Figure S3.

As we did not find substantial doxycycline-induced transcriptional changes that could sufficiently explain the protective effects of the drug, we then investigated the liver metabolic profiles in the presence of doxycycline and infection, given the centrality of liver function and metabolic changes during sepsis (Fleury et al., 2019; Ganeshan et al., 2019). To this end, we used an untargeted metabolomics approach to study metabolic changes in the liver 8h after infection. Analysis of the top up- and down-regulated metabolites identified two main signatures that identified a pronounced accumulation of acylcarnitines and glucocorticoids in the liver of infected mice (Figure 3C). While glucocorticoids have an important role in modulating both inflammation and metabolism during infection, the abnormal acylcarnitine profile points to a defective import and oxidation of fatty acids into the mitochondria, which may be a cause of liver and metabolic failure during sepsis. To further validate and investigate the identified FAO signature, we then measured mRNA levels of *Pparα*, a known master regulator of fatty acid oxidation (FAO), and several of its transcriptional targets, including *Cpt1*, *Cpt2* and *Slc25a20*, responsible for free fatty acid (FFA) import into the mitochondria, at 8 and 24h after infection. We consistently found a decreased expression of all of the analyzed targets upon infection, which was maintained at least for the first 24h (Figure 3D), thus highlighting lipid metabolic dysfunction as a hallmark of sepsis pathophysiology. To functionally and causally test the importance of FAO for the sepsis progression and survival, we treated infected mice with etomoxir (Divakaruni et al., 2018), a frequently used inhibitor of CPT1a – the enzyme that catalyzes the conversion of free fatty acids to acylcarnitines before their import into the mitochondrial matrix. We observed a significant increase in mortality as compared to infected non-treated controls (Figure 3E). This result was substantiated by liver specific shRNA-mediated silencing of *Cpt2*, another key component of long chain fatty acid import into the mitochondria (Figure S3C and D). A similar effect was found for glucocorticoid response, as mice treated with mifepristone, a glucocorticoid receptor (GR) antagonist, had significantly worse infection outcomes (Figure 3F). Taken together, these results demonstrate that both FAO and response to glucocorticoids are necessary for recovery from sepsis, and that impairment of these pathways correlates with worse infection outcomes.

### Doxycycline improves FAO and response to glucocorticoids

Having identified FAO and response to glucocorticoids as necessary for survival in sepsis, we next investigated the effect of doxycycline these pathways. Considering the lack of transcriptional signatures in mouse liver upon doxycycline treatment (Figure 3B), we turned to an HPLC-MS analysis to identify several acylcarnitine and FFA species in mouse liver 8h after infection with and without doxycycline treatment. We found that doxycycline partially corrects the accumulation of acylcarnitines and FFA upon infection (Figure 4A), suggesting that it may improve the block in fatty acid transport into the mitochondria caused by infection. Remarkably, several attempts to correct FFA transport in the liver by overexpressing several genes involved in this pathway, alone or in combination, using an adeno-associated virus (AAV) serotype 8 driven by liver thyroid hormone-binding globulin (TBG) promoter, conferred no survival advantage during infection (Figure S4A, S4B, S4C). An attempt to bypass the carnitine shuttle by orally supplementing mice with octanoic acid, a medium-chain FFA (C8:0) that does not require active transport into the mitochondria or conjugation with carnitine, was also not capable of consistently rescuing mice (Figure S4D).

**Figure 4.**
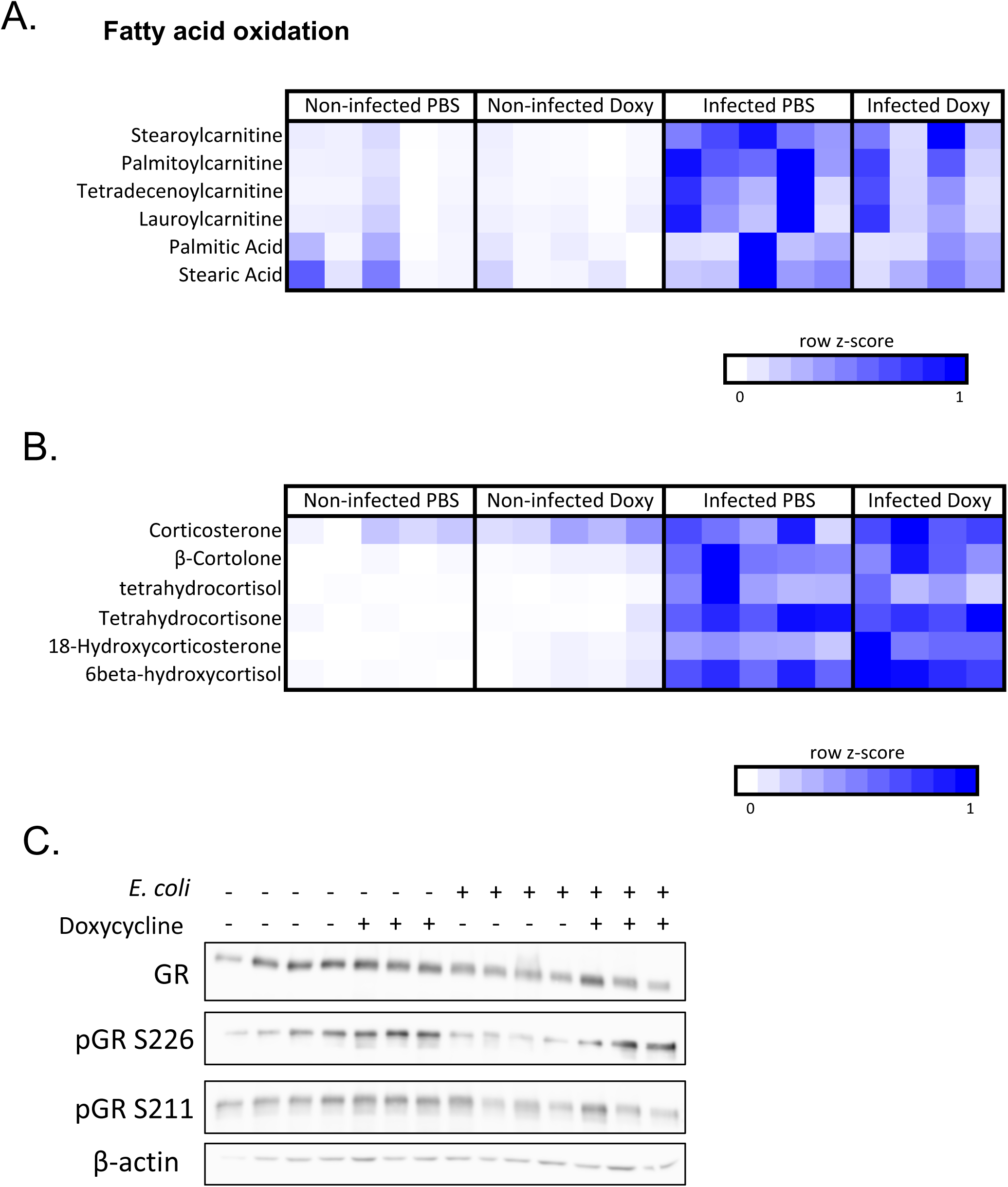
Doxycycline improves both FAO and response to glucocorticoids. (A), HPLC-MS analysis of FAO metabolites in mouse liver 8h after infection and/or doxycycline treatment. Each square represents one mouse. (B) Glucocorticoid levels from an untargeted metabolomics analysis in mouse liver 8h after infection and/or doxycycline treatment. Each square represents one mouse. (C) Protein levels of total and phosphorylated glucocorticoid receptor (GR) in mouse liver at 8h after infection or doxycycline treatment. Each lane represents one mouse. See also Figure S4.

Likewise, treatment with the PPARα agonist CP868388 at the time of infection was not enough to improve survival (Figure S4E). Levels of glucocorticoids were sharply elevated in the liver in response to infection (Figure 4B). The effect of doxycycline in the levels of glucocorticoids was marginal in the cases of infected and non-infected mice (Figure 4B). Sepsis is characterized by resistance to glucocorticoids, which means that even high levels of these species, both endogenously produced and therapeutically administered, fail to produce their anti-inflammatory and metabolic modulator effects (Dendoncker and Libert, 2017). In fact, pre-treatment of mice with the synthetic glucocorticoid dexamethasone before infection with *E. coli* failed to significantly increase survival (Figure S4F). To explore glucocorticoid signaling in this context, we probed liver extracts collected 8h after infection with and without doxycycline treatment for total glucocorticoid receptor (GR) and two of its phosphorylated forms, on Ser226 and Ser211, which are markers of GR activation (Bouazza et al., 2012). Infected, PBS-treated mice showed markedly reduced levels of all analyzed forms, supporting the hypothesis of glucocorticoid resistance in sepsis (Figure 4C). Interestingly, doxycycline treatment increases phospho-Ser226 and to a lesser extent phospho-Ser211, both in the presence and absence of infection, while total GR levels are also moderately increased in doxycycline-treated, *E. coli* infected mice (Figure 4C). These observations indicate that doxycycline does not significantly affect the levels of endogenous glucocorticoids in infected or non-infected conditions, but substantially increases the activation of the GR in response to glucocorticoids, which is normally blunted in sepsis (Dendoncker et al., 2019). Together, these results suggest that, while FAO and glucocorticoid signaling are perturbed and are necessary for survival in sepsis, independently rescuing these pathways is not sufficient to substantially improve the outcome of a severe infection. Notably, doxycycline treatment is able to improve both FAO and glucocorticoid signaling, which may contribute substantially for tissue function maintenance and metabolism during sepsis.

### Low-dose doxycycline affects mitochondrial function in vivo

Having phenotypically characterized the salutary effects of doxycycline for survival to sepsis, tissue homeostasis, and metabolism, we then turned to the mechanistic identification doxycycline-induced perturbations leading to disease tolerance. The role of microbiota in host physiology has been increasingly acknowledged, with extensive research focusing on its therapeutic potential in sepsis (Haak et al., 2018) including its role in disease tolerance (Schieber et al., 2015). As an antibiotic, doxycycline induces substantial changes in gut microbiome composition (data not shown) that may indirectly affect host fitness. To address role of the contribution of microbiota for doxycycline-induced protection against sepsis, we tested the effect of doxycycline treatment in the TetR *E. coli* sepsis model in C57BL/6J mice raised and maintained in germ-free conditions. Both survival (Figure 5A) and body temperature (Figure S5A) were significantly improved in doxycycline-treated mice whereas no differences were found in body weight (Figure S5B). These results largely phenocopy the protective effect observed on conventionally raised, specific pathogen-free mice (Figure 1A, 1B, 1C), demonstrating a host-dependent disease tolerance mechanism.

**Figure 5.**
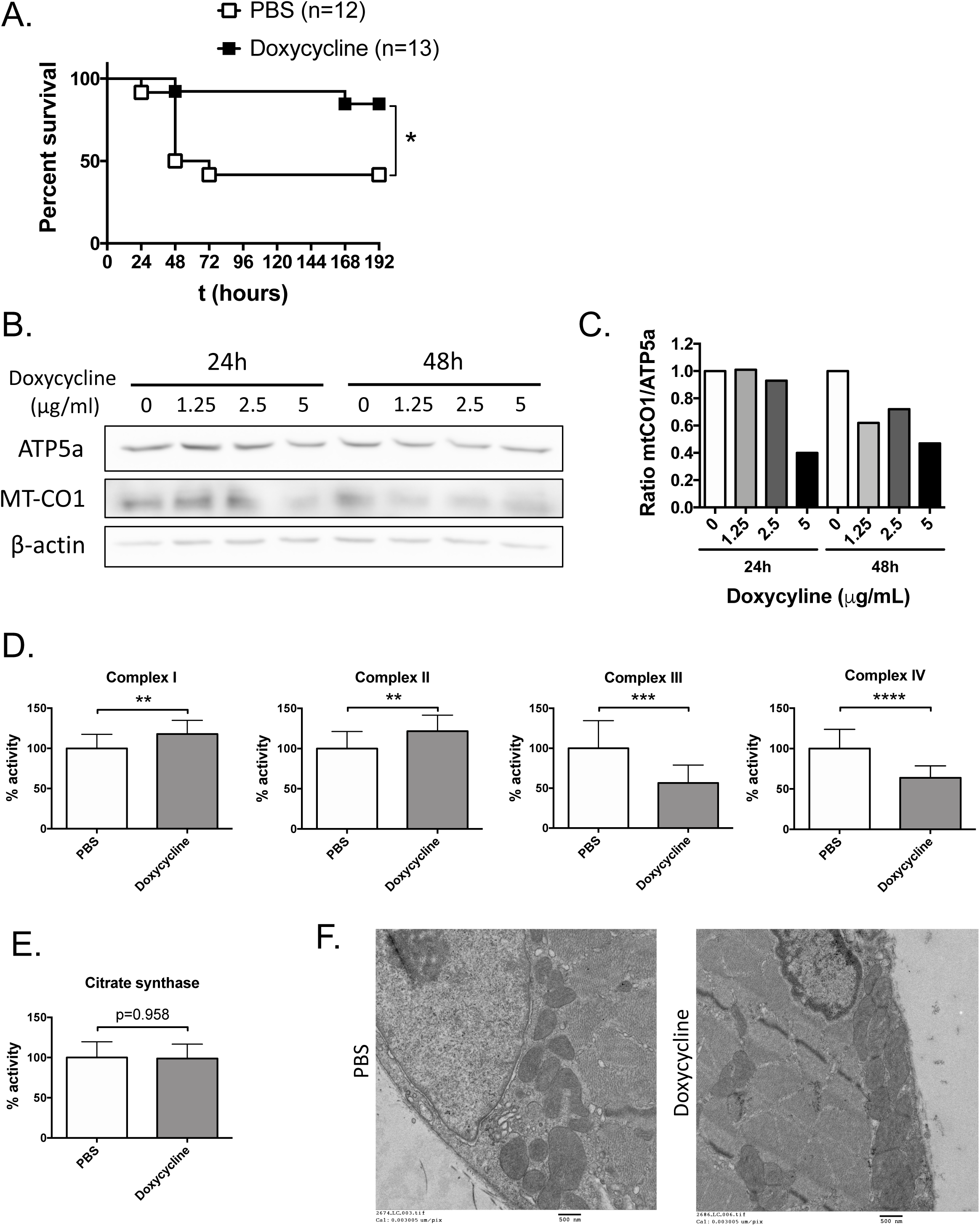
Low-dose doxycycline affects mitochondrial function *in vivo.* (A) Survival after infection of germ-free mice with TetR CamR *E. coli* and treatment with 1.75 μg/g body weight doxycycline (or PBS as a control) at 0, 24 and 48 h. (B) Immunoblot of HepG2 cells incubated with varying concentrations of doxycycline for 24 or 48h and probed for nuclear (APT5a) or mitochondrial (MTCO1)-encoded proteins of the ETC. (C) MT-CO1/ATP5a ratio determined by densitometry in (B). (D) Enzymatic activity of the ETC complexes I, II, III, and IV in mouse liver collected 12h after PBS or doxycycline treatment. Enzymatic activity is expressed as % of PBS-treated control, normalized for citrate synthase (CS) activity. (E) CS activity (expressed as % of control) in mouse liver collected 12h after PBS or doxycycline treatment. (F) Representative images of transmission electron microscopy in mouse skeletal muscle 24h after PBS or doxycycline treatment. Scale bar = 500 nm. (A) Represents pooled data from four independent experiments. Data in (D) and (E) represent mean±SD of 12 mice/group from two independent experiments. See also Figure S5.

Having documented a direct host-dependent effect of doxycycline leading to disease tolerance against sepsis, we then explored the hypothesis that doxycycline protection is initiated and dependent on mitochondrial function perturbations. We first explored the possibility that doxycycline treatment could increase the availability of the elongation factor Tu (TUFM) as this antibiotic works by blocking the binding of aminoacyl-tRNA (aa-tRNA) to the A site of the ribosome, decreasing the use of TUFM which is part of the ternary complex (aa-tRNA, TUFM and GTP) that decodes the gene open reading frame (Lin et al., 2018). If this would be the case, increased availability of TUFM could play a role in disease tolerance because TUFM has been shown to interact with NLRX1 causing reduction of type I interferon and enhancement of autophagy (Lei et al., 2013). To test this hypothesis, we again used the adeno-associated virus (AAV) serotype 8 driven by liver thyroid hormone-binding globulin (TBG) promoter system to overexpress TUFM in the liver. While the overexpression of TUFM in the liver was effective (Figure S5C, right panel), we did not observe a substantial improvement in survival (Figure S5C, left panel) and therefore conclude that this mechanism does not play a substantial role in the promotion of disease tolerance induced by doxycycline against severe bacterial infection.

Doxycycline has been reported to block synthesis of mitochondrial-encoded proteins at relatively high doses across several model organisms, including mammalian cells (Michel et al., 2015; Moullan et al., 2015). We used the human hepatocellular carcinoma cell line HepG2 to examine the relative abundance of proteins of the ETC encoded by the nucleus (ATP5a) and by the mitochondrial DNA (MT-CO1) upon incubation with doxycycline. Even low concentrations of the drug (1.25 to 5 μg/mL) showed decreased MT-CO1 levels while ATP5a levels remain unchanged (Figure 5B, 5C), demonstrating that pharmacologically relevant doses of doxycycline have an impact in mitochondria. Remarkably, functional changes in mitochondrial function could also be detected *in vivo*, by measuring the enzymatic activity of the ETC complexes I, II, III, and IV in mouse liver collected 12 h after doxycycline treatment. We found strongly reduced activity of complexes III and IV (54% and 60% of the control, respectively) and a slight, possibly compensatory, activation of complexes I and II (115% and 118% of the control, respectively) (Figure 5D). These changes in function were, however, not reflected in impaired mitochondrial integrity as judged by citrate synthase (CS) activity (Figure 5E), or morphology as analyzed by transmission electron microscopy (Figure 5F). Inhibition of ETC function can lead to the generation of reactive oxygen species (ROS) (Zhao et al., 2019). To investigate the effect of doxycycline treatment in the generation of ROS we treated HepG2 cells with doxycycline for 24 h and used the MitoSOX fluorescent reporter to measure peroxide ROS generation by cytometry. A low dose of doxycycline, 1.25 and 5 μg/mL, induced a modest increase in ROS levels in HepG2 cells (Figure S5D), without affecting the mitochondrial mass as revealed by the use of Mitotracker green (Figure S5E). The ATM kinase, that we previously have shown to be required for the activation of disease tolerance in response to low dose anthracycline treatment (Figueiredo et al., 2013) can be activated by ROS (Choy and Watters, 2018). To test the hypothesis that doxycycline induction of disease tolerance could be mediated by the ROS activation of ATM, we have used AAV8-CRE mediated depletion of ATM in the liver and monitored survival in response to infection. We found that liver-depletion of ATM did not substantially affect the protective phenotype of doxycycline (Figure S5F) and conclude that the doxycycline induction of disease tolerance is independent of ATM activation in clear contrast to the case of anthracyclines (Figueiredo et al., 2013). Seahorse based measurement of oxygen consumption rate in response to doxycycline in A549 cells showed only a marginal decrease in mitochondrial basal respiration and maximal respiratory capacity (data not shown) in good agreement with previously published results (Kalghatgi et al., 2013). Taken together, our data indicate that low-dose doxycycline triggers changes in mitochondrial ETC function without compromising mitochondrial integrity or other major mitochondrial functional features, which may explain the observed changes in liver mitochondrial metabolism (Figure 4A).

### Mild transient perturbations in mitochondrial ETC increase survival in sepsis

To determine a causal link between mild perturbations in mitochondrial ETC function and disease tolerance, we next used a genetic system to induce such perturbations. CRIF1 is a mitoribosomal protein with an important role in the assembly and function of ETC complexes (Kim et al., 2012). A previous study linked tissue-specific *Crif1* knockout with reduced ETC activity, resulting in systemic metabolic benefits (Chung et al., 2017). Prompted by the doxycycline-induced improvement in liver pathology (Figure 1E, 1F, 1G) and metabolism (Figure 4A) during sepsis, we decided to test the impact of targeted liver deletion of CRIF1 in sepsis. To this end, we took advantage of the AAV8-TBG strategy to obtain fast, efficient and specific gene editing in mouse liver. Intravenous injection of Cre-expressing AAV8 in homozygous *Crif1^lox/lox^* or heterozygous *Crif1*^lox/-^ mice led to high protein levels of Cre recombinase after 7 days, accompanied by reduced CRIF1 protein and mRNA levels (Figure 6A). Mice homozygous for the flox allele suffered almost full deletion in mRNA (98%) and protein levels, whereas heterozygous mice showed a 30% decrease in mRNA and only modest decrease in protein levels (Figure 6A). Strikingly, *E. coli* infection performed 7 days after AAV8-Cre injection resulted in increased survival of *Crif1*^lox/-^ mice but not *Crif1^lox/lox^* littermates (Figure 6B), thus supporting the notion of a beneficial role for mild, transient mitochondrial perturbations but deleterious effect for severe perturbations of ETC mitochondrial function. *Crif1*^lox/-^ mice also showed less severe hypothermia (Figure S6A), similar to doxycycline-treated mice (Figure 1B), but no differences in body weight when compared to *Crif1^lox/lox^* mice and to mice injected with a control AAV-GFP vector (Figure S6B). Bacterial loads in blood of *Crif1*-depleted mice were similar to the controls (Figure 6C), supporting the notion of disease tolerance induced my mild mitochondrial perturbations.

**Figure 6.**
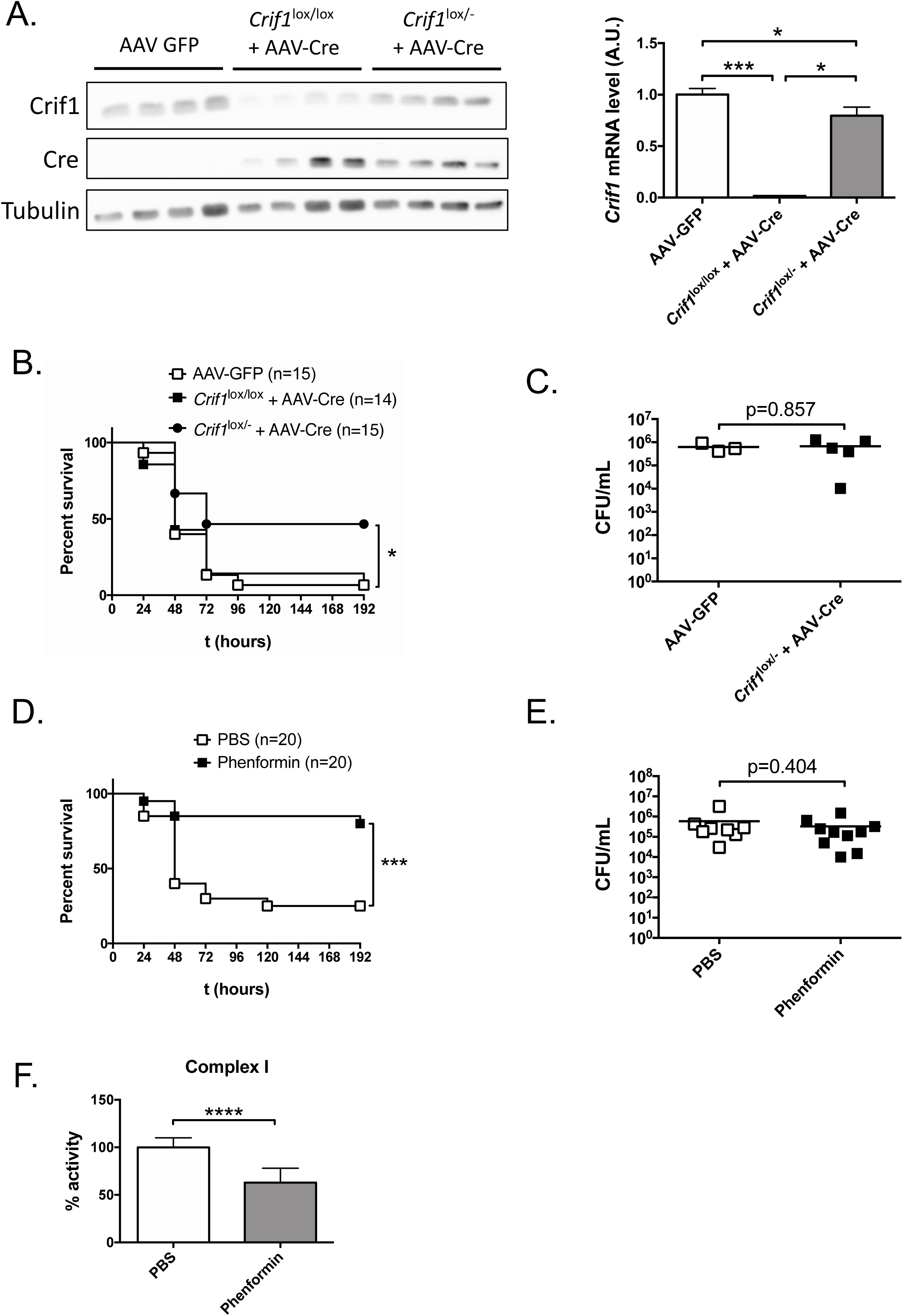
Mild, transient perturbations in mitochondrial function are associated with increased survival in sepsis. (A) Crif1 and Cre recombinase protein levels and *Crif1* mRNA levels in *Crif1*^lox/lox^ or *Crif1* ^lox/-^ mice 7 days after injection of liver-specific AAV8-TBG expressing Cre recombinase (or GFP as a control). (B) Survival after infection of *Crif1*^lox/lox^ or *Crif1* ^lox/-^ mice with TetR CamR *E. coli*. Mice were previously injected with AAV8-TBG expressing Cre recombinase (or GFP as a control). (C) Bacterial loads in *Crif1* ^lox/-^ mouse blood 24h after infection. (D) Survival after infection of C57BL/6J mice with *E. coli* and treatment with 100 μg/g body weight phenformin (or PBS as a control) at the time of infection. (E) Bacterial loads in blood of PBS and phenformin-treated mice, 24 h after *E. coli* infection. (F) Enzymatic activity of the ETC complex I in mouse liver collected 12h after PBS or phenformin treatment. ETC activity is expressed as % of PBS-treated control, normalized for CS activity. Data in (A) represents mean±SD of 4 mice/group assayed in triplicate. (B) Represents pooled data from three independent experiments. (C) Represents data from a single experiment. Bars indicate the mean. (D) Represents pooled data from four independent experiments. (E) Represents pooled data from two independent experiments. Bars indicate the mean. (F) Represents mean±SD of 5 mice/group from a single experiment. See also Figure S6.

To address the question of whether the induction of disease tolerance by ETC required general or specific perturbations of ETC complexes, we next focused on the biguanide antidiabetic drug phenformin, previously reported to specifically and potently inhibit complex I of the ETC (Dykens et al., 2008). We found that injection of a single dose of phenformin at the time of *E. coli* infection resulted in remarkable survival advantage (Figure 6D) and improvement in body temperature control (Figure S6C), with modest effects in body weight (Figure S6D). Interestingly, administration of two doses of phenformin, separated by 24 h, lead to increase mortality in striking contrast to a single administration (Figure S6E). Similar to doxycycline, phenformin confers protection to sepsis by inducing disease tolerance mechanisms, as bacterial load in mouse blood collected 24h after infection shows no difference between PBS and phenformin-treated mice (Figure 6E). We then evaluated the enzymatic activity of ETC complexes in mouse liver collected 12h after phenformin treatment and concluded that complex I activity in dramatically decreased (61% of the control) (Figure 6F), while complexes II, III and IV, as well as CS show little to no changes (Figure S6F). Additionally, phenformin-treated mice present lower levels of serum inflammatory markers (Figure S6G) and lower scores of histological damage (Figure S6H). In summary, we show that mild and transient perturbations in mitochondrial function, which may affect the activity of different complexes of the ETC, activate disease tolerance mechanisms in a mouse model of bacterial sepsis.

## Discussion

Both resistance and disease tolerance mechanisms are necessary to minimize loss of function in response to infection and survival (Soares et al., 2017). Resistance mechanisms have been extensively explored, but those responsible for disease tolerance are for the most part not understood. The requirement for synergy of both defense strategies is perhaps best illustrated in sepsis where effective control and elimination of the inciting pathogen is not sufficient to ensure survival, especially in cases where tissue damage and loss of function are pronounced. We have previously shown that DNA damage responses initiated by low dose anthracyclines can robustly protect in several mouse models of sepsis, by activating disease tolerance mechanisms in an ATM- and autophagy-dependent manner (Figueiredo et al., 2013). Here we asked whether perturbation of other core cellular functions could also lead to protective responses against severe infection. Using an *in vivo* small chemical screen approach, we identified tetracycline antibiotics as a novel class of drugs capable of activating disease tolerance mechanisms. Our findings are in good agreement with the long-known effects of several classes of antibiotics on the course and outcome of infections that cannot be explained by their direct antibacterial activities alone (Tauber and Nau, 2008). For example, macrolides – a class of lactone ring antibiotics – have extensively documented clinical roles in chronic inflammatory pulmonary disorders (Rubin and Henke, 2004; Sharma et al., 2007), and at least experimentally, members of this class like erythromycin, have also demonstrated protective effects several models of cerebral ischemia (Brambrink et al., 2006). Other classes, including fluoroquinolones and tetracyclines (the focus of the current work), have also been labeled as immunomodulators but their effects in addition to their antibiotics properties have not been described beyond the phenotypic modulation of inflammatory pathways, for which the molecular mechanistic bases remain unidentified (Tauber and Nau, 2008). Here we show that the strong protective effects of tetracycline antibiotic doxycycline against sepsis relies on host-dependent induction of disease tolerance. Mechanistically, disease tolerance is initiated by the inhibition of mitochondrial protein synthesis which leads to the causal perturbation of ETC function.

Our results show that disease tolerance induced by doxycycline are pathogen group specific as survival is robustly enhanced in a bacterial model of sepsis, the speed of recovery is increased in a model of viral lung infection, but no salutary effects were observed in the case of a fungal infection (*C. albicans*) or the rate of cerebral malaria in spite of the substantial decrease in parasitemia, as it would be expected by the sensitivity of plasmodium species to tetracycline antibiotics (Pagès et al., 2002). These findings support the notion that tissue damage control mechanisms are likely to function in pathogen-specific manner to confer disease tolerance to different types of infection because they impose distinct forms of stress and damage to their hosts that require dedicated tissue repair responses (Soares et al., 2017; Wang et al., 2016).

To specifically study the disease tolerance effects of doxycycline, we used a model of sepsis that initiates infection with an i.p. injection of an *E. coli* that was made resistant to tetracyclines (TetR) to control for the effects of this antibiotic directly on bacterial viability (resistance). Even in this model, it would remain possible that the protective effects of doxycycline (or other ribosome-targeting antibiotics) could be due to its effects on the microbiota, which could shift towards a more tolerogenic, pro-repair profile. To test this possibility, we administered doxycycline in a similar model of sepsis, but using germ-free C57BL/6J instead of the regular SPF C57BL/6J mice. As germ-free mice can still be robustly protected by doxycycline against a TetR *E. coli* (Figure 5A), we conclude that regardless of a potential protection through the modulation of the microbiome, ribosome-targeting antibiotics like tetracyclines induce disease tolerance leading to protection against infection by affecting host processes. This protection relies on a better tissue damage control as demonstrated by lower release of serologic markers of organ lesion (Figure 1E) and substantially improved histopathology that was especially striking in the liver and lung (Figure 1F). While we have also observed significant decreases of critical inflammatory markers of sepsis in doxycycline-treated mice, including TNFα and IL-6 (Figure 1H), these effects are likely indirect and reflect systemic improvement and progress of the infection, because *in vitro* induction of these mediators is not modified by a wide range of doxycycline concentration treatments (Figure S1D).

By further charactering the effects of doxycycline in the organs where tissue damage was most improved at the histopathological level, we found important reasons for improved survival of treated mice. In the lung, comparing global transcriptional profiles of doxycycline-treated and non-treated mice, both in infected and non-infected conditions identified a signature that suggests an increased repair response, possibly by recruiting progenitors and inducing their differentiation to substitute for the infection-damaged epithelial cells. Our data suggest that doxycycline represses the expression of stem cell markers, providing a signal for differentiation, which may result in improved lung repair upon infection. Some of these cell populations are rare and not well characterized but seem to have an important role in tissue repair mechanisms (Xian and McKeon, 2012). While the transcriptional signature points to basal stem cells, its partial genetic ablation does not significantly impair the survival of infected mice (Figure S2D) raising the possibility that doxycycline recruits additional progenitor populations to account for the complete repair response. The identity and signaling events in these populations induced by doxycycline is currently under investigation, but it is possible that doxycycline activates lung cell progenitors by affecting their oxphos/glycolysis balance as demonstrated previously for stem cells (Chang et al., 2014; Qureshi et al., 2017).

Surprisingly, when investigating more deeply the effects of doxycycline in the liver, we did not observe the same transcriptional signature present in the lung. In fact, we found almost no significant transcriptional changes induced by doxycycline regardless of the infection state of the animals. We therefore used other unbiased technologies to identify possible causal changes leading to improvements of infection in treated animals. Using untargeted metabolomics, we identified two clear molecular signatures induced by infection in the liver. One points to the clear accumulation of acylcarnitines and FFA upon infection (Figure 4A) and suggests that severe infection induced a block in long chain fatty acid transport and oxidation in the mitochondria (Figure 3C). In fact, the key mediators of this transport (*Cpt1a*, *Cpt2* and *Slc25a20*) and their master transcriptional regulator (*PPARα*), are all strongly repressed by infection, in good agreement with recent observations of LPS suppressed program of fatty acid oxidation (Ganeshan et al., 2019) and the requirement of liver PPAR*α* for metabolic adaptation to sepsis (Paumelle et al., 2019). The other signature we found points to a sharp increase in multiple glucocorticoid species (Figure 3C), suggesting that the infection imposes a strong stress response in the liver, but also to one of the known hallmarks of sepsis pathology: resistance to glucocorticoid treatment (Dendoncker et al., 2019; Van Wyngene et al., 2018). We do find that infection decreases markers of GR activation like the phosphorylation of Ser211 and Ser226 (Figure 4C). Both of these signatures that are severely repressed by infection are critical for survival as demonstrated decreased survival of infected animals by etomoxir-mediated inhibition of CPT1 (Figure 3E) and GR inhibition by mifepristone (Figure 3F). Interestingly, modulation of each of these processes alone is not enough to substantially improve survival (Figure S4), but doxycycline positively impacts on both of these processes (Figure 4), which likely contributes decisively to the robust increase in survival by a favorable metabolic reprograming in the liver in response to a severe infection. In fact, this shift to catabolic programs in parenchymal tissues has been associated with tissue protection, resistance to stress, and disease tolerance across several experimental models (Medzhitov, 2019).

The positive effects induced by doxycycline treatment, both in the lung and in the liver, are likely to be initiated by the partial and transient perturbation of ETC activity that we document in this study, pharmacologically and genetically (Figure 6). This effect may be due to the observed mito-nuclear imbalance caused by inhibition of protein synthesis in the mitochondria (Figure 5B and C) that cause most prominent effects in the activities of ETC complexes III and IV (Figure 5D). Interestingly, gaso-transmitters like carbon monoxide, that is known to inhibit complex IV activity (Zuckerbraun et al., 2007), has been reported to induce disease tolerance and promote survival to malaria (Pamplona et al., 2007). Based on these facts, we asked whether the protective effects of ETC function perturbation were specifically due to complex IV inhibition or perturbation of the activity of other complexes could also induce disease tolerance. The perturbation of any of the ETC complexes alone seems to be able to induce disease tolerance to infection as phenformin, a strong complex I inhibitor (Figure 6F), robustly protects from sepsis by inducing disease tolerance, not resistance mechanisms (Figure 6D and 6E).

Our data suggests that ETC perturbations triggers not only local protective responses but also systemic effects, as demonstrated by improved survival by direct administration of doxycycline to the lung (Figure 2A) and by liver specific genetic deletion of CRIF1 (Figure 6B). Both in the case of ETC function perturbations induced genetically and pharmacologically, the protection can be induced by partial but not very strong perturbations. These findings suggest, as we have shown previously in the case of DNA damage induced by low dose anthracyclines (Figueiredo et al., 2013), an hormetic response induced by perturbations of core cellular functions that induced compensatory cytoprotective responses against infection (Medzhitov, 2013). In addition to the protective role of ATM-dependent DNA damage responses, our current study adds a novel pathway leading to disease tolerance that can be initiated by inhibition of mitochondrial protein synthesis leading to perturbations of ETC function. The elucidation of the signal generated by ETC function perturbation, the mechanisms that sense it and its downstream transduction and molecular effectors are challenging but critical goals to understand cytoprotection and disease tolerance, which ultimately can be key for the development of novel therapies against infection, sepsis in particular.

## Author contributions

Conceptualization, LFM with input from HGC; Methodology, HGC, AB, VB and TK; Formal analysis, HGC, AB, VB and TK; Investigation, HGC, AB, ANC, ES, DP, TV, KW, VB, SW and TK; Resources, HSY and MS; Data analysis and curation, AB; Writing - original draft, HGC and LFM; Supervision, LFM; Funding acquisition, LFM.

## Acknowledgements

The authors would like to acknowledge M.J. Amorim (IGC) for the influenza virus, MP. Soares and S. Ramos (IGC) for assistance with *Plasmodium* infection, S. LeibundGut-Landmann (U. Zurich) for the *C. albicans* strain, RB. Soria and I. Gordo (IGC) for TetR phage lysate, and W. Xian and F. McKeon (Harvard Medical School, Boston, MA, USA) for the KRT6-DTR mice. We are grateful for the technical support from the Animal House, Histopathology Unit, Electron Microscopy Facility, and Genomics Unit at IGC. This work received financial support from the European Community Horizon 2020 (ERC-2014-CoG 647888-iPROTECTION) and Fundação para a Ciência e Tecnologia (FCT: PTDC/BIM-MEC/4665/2014). HCG is recipient of an FCT fellowship (PD/BD/105998/2014). The VBCF Metabolomics Facility is funded by the City of Vienna through the Vienna Business Agency.

## Lead Contact and Materials Availability

Further information and requests for reagents may be directed to and will be fulfilled by the Lead Contact Luís F. Moita. This study did not generate new unique reagents, except for the TetR CamR *E. coli* strain.

## Experimental Model and Subject Details

### Mice

All animal studies were performed in accordance with Portuguese regulations and approved by the Instituto Gulbenkian de Ciência ethics committee (reference A002.2015) and DGAV. C57BL/6J mice were obtained from Instituto Gulbenkian de Ciência or Charles River Laboratories (France). Male mice, 8 to 12 weeks old were used, except if otherwise stated. *Crif1*^lox/lox^ mice (Kwon et al., 2008) were obtained from M. Shong (Chungnam National University School of Medicine, Daejeon, South Korea). KRT6-DTR mice (Zuo et al., 2015) were obtained from W. Xian and F. McKeon (Harvard Medical School, Boston, MA, USA). *Atm*^-/-^ mice (Borghesani, 2000) were obtained from F. Alt (Harvard Medical School, Boston, MA, USA). Mice were maintained under specific pathogen-free (SPF) or germ-free (GF) conditions with 12 h light/12 h dark cycle, humidity 50–60%, ambient temperature 22 ± 2°C and food and water *ad libitum*. For all experiments, age-matched mice were randomly assigned to experimental groups.

### Cell lines

HepG2 (male) human hepatocellular carcinoma cells were cultured in DMEM supplemented with 10% (v/v) FBS and 1% (v/v) Penicillin-Streptomycin at 37°C with 5% CO_2_. Three to five days before experiments, medium was changed to DMEM with 10% FBS with no addition of antibiotics.

### Primary cell cultures

BMDMs were differentiated from adult (typically 8-week-old) C57BL/6J male mice. After euthanasia by CO_2_ inhalation, the mouse skin was sterilized with 70% ethanol and femurs and tibia of hind limbs were removed, stripped of muscle and rinsed in RPMI medium. Bone marrow cells were flushed from cut bones using an insulin syringe with a 30G needle into 10 mL of RPMI medium. Cells were then pelleted by centrifugation at 450 g for 5 min and the cell pellet resuspended in 10 mL of RPMI supplemented with 10% (v/v) FBS and 0.2 % (v/v) penicillin/streptomycin. Cells were counted and plated at a density of 3 x 10^6^ cells (including red blood cells) per 10 mL of RPMI medium supplemented with 10% FBS and 0.2 % penicillin/streptomycin, with 30% of L929-conditioned medium. After three days, an equal volume of fresh medium with 30% of L929-supernatant was added to the cells. After four additional incubation days, the medium was replaced by 10 mL of fresh medium with 30% of L929-supernatant. 24 h-48h afterwards, cells were scraped from plates, counted and seeded in C10 medium. L929-conditioned medium: L929 cells were cultured in T175 flasks, in 40 mL of DMEM medium with 10% (v/v) FBS and 1% (v/v) Penicillin/Streptomycin and grown to confluency. The culture medium was left unchanged for 5 days, for good production of M-CSF. Cells were then centrifuged at 290 g for 5 minutes and the supernatant was collected and filter sterilized. C10: RPMI medium 1640 supplemented with: 10% (v/v) Fetal Bovine Serum (FBS), 1% (v/v) Penicillin-Streptomycin, 1% (v/v) Pyruvate, 1% (v/v) LGlutamine, 1% (v/v) Non-essential aminoacids, 1% (v/v) Hepes buffer, 0.05 M of 2-Mercaptoethanol.

### Bacterial cultures

*Escherichia coli* K12 MG1655 carrying resistance to chloramphenicol was made resistant to tetracyclines by P1 phage transduction (P1 phage lysate was a gift of Roberto Balbontín from the Evolutionary Biology group at the IGC). All bacterial cultures were carried out in Luria-Bertani broth supplemented with 10 μg/mL doxycycline (LB+doxy), except for survival studies in chloramphenicol-treated mice, in which bacterial cultures were made in LB + 50 μg/mL chloramphenicol.

### Fungal cultures

*Candida albicans* (Robin) Berkhout (Gillum et al., 1984) were cultured in yeast culture medium (YPD) for 16-20h at 30°C, 180 rpm.

## Method details

### E. coli-induced sepsis model and drugs treatments

A starter culture from a single *E. coli* colony was incubated overnight (12-16h) at 37°C, 200 rpm. The next morning, the culture was diluted 1:50 in LB+doxy and incubated for 2.5 hours until late exponential phase was reached (OD600nm = 0.8-1.0). The culture was then centrifuged at 4400 x*g* for 5 minutes at room temperature, washed with PBS and resuspended in PBS to obtain an OD_600nm_ = 4.5-5.0, corresponding to 1-2×10^9^ CFU/mL. This bacterial suspension was immediately injected intraperitoneally (200 μL/mouse) in mice using a 27G-needle. Infections were always performed in the morning. The concentration of the inoculum was determined by plating 10^-6^ and 10^-7^ dilutions in LB+doxy agar plates and incubating overnight at 37°C. Doxycycline hyclate was dissolved in PBS and injected intraperitoneally (200 μL/mouse) at 1.75 μg/g body weight 0, 24 and 48h after infection. The following drugs were dissolved in the indicated vehicles and injected intraperitoneally (200 μL/mouse, except mifepristone: 50 μL/mouse) at the time of infection and at the indicated concentrations: chloramphenicol (vehicle: 5% cyclodextrin, dose 50 μg/g); phenformin (vehicle: PBS, dose 100 μg/g); etomoxir (vehicle: PBS, dose 15 μg/g); mifepristone (vehicle: 100% DMSO, dose 30 μg/g); CP868388 (vehicle: 7% DMSO, dose 3 μg/g), dexamethasone (vehicle: PBS, dose 5 μg/g). Octanoic acid was dispersed in 0.5% methylcellulose and supplemented by oral gavage (200 μL/mouse) at 2, 8, 24, and 48h post-infection. Intra-tracheal administration of doxycycline was performed as previously described (DuPage et al., 2009). Briefly, mice were anesthetized with an intraperitoneal injection of 450 μg/g avertin, placed on an intubation platform (Labinventions.com), and intubated using a 22G, 1-inch catheter. Doxycycline (1.75 μg/g body weight in 50 μL PBS) was then pipetted into the opening of the catheter and the catheter was removed after all volume was inhaled. Mice were allowed to recover from anesthesia and infection with *E. coli* was performed two hours later. Body weight and rectal temperature were determined 0, 24 and 48h after infection. For survival experiments, mice were closely monitored during one week for survival and health status. Moribund animals (i.e. shivering or unable to maintain upright position) were euthanized. For tissue analysis, mice were sacrificed at the indicated time-points by CO_2_ inhalation, blood was collected by cardiac puncture and organs were harvested, immediately frozen in liquid nitrogen and stored at −80°C. Blood was centrifuged at 1600 x*g* for 5 min and serum collected and stored at - 80°C.

### Other infection models

For influenza virus infection, mice were anesthetized by inhalation of isoflurane and intranasally inoculated with a sublethal (100 pfu/mouse) dose of Influenza A PR/8 (Wit et al., 2004) in 30 μL PBS. Infected mice were treated with an intraperitoneal injection of 1.75 μg/g doxycycline at days 4, 5 and 6 post-infection. Infection with GFP-transgenic *Plasmodium berghei* ANKA was performed as described (Pamplona et al., 2007). Briefly, female mice 8-12 weeks old were given an intraperitoneal injection containing 1×10^5^ infected red blood cells from a previously infected mouse. Doxycycline (1.75 μg/g body weight) was injected daily starting at the time of infection. Blood samples were taken from the tail vein and analyzed in FACSCalibur to determine parasitemia (expressed as % of GFP-positive red blood cells). *C. albicans* cultures were washed and resuspended in PBS to obtain an OD_600nm_ = 0.5, corresponding to 5×10^6^ CFU/mL. Female mice 8-12 weeks old were infected by an intravenous injection of 100 μL in the tail vein. Mice were treated with 1.75 μg/g body weight doxycycline at 0, 24 and 48h after infection.

### Liver-specific gene editing with Adeno-associated virus (AAV)

AAV serotype 8 constructs were diluted in sterile PBS and 5×10^11^ gc/mouse were delivered by retro-orbital injection. All subsequent experiments were performed 7 days after AAV injection.

### Ablation of KRT6^+^ cells

Mice hemizygous for the human diphtheria toxin receptor inserted in the *Krt6a* locus (Krt6a-DTR) and wild-type littermates were treated with 12 ng/g diphtheria toxin (DT) by intra-tracheal administration, as described above. Experiments were performed 4 days after DT administration.

### Colony Forming Units assay

Freshly collected samples of liver, lung and kidney were homogenized in 1 mL sterile PBS using TissueLyser II (Qiagen). Colony forming units (CFU) were determined in blood and organs by serially diluting in sterile PBS and plating in LB+doxy agar plates. At least three dilutions were plated per condition. CFU were counted after incubating plates at 37 °C for 16h.

### Biochemical assays in mouse serum and supernatant from BMDMs

Cytokine levels were determined using the ELISA kits indicated in the KRT, according to the manufacturer’s instructions. Serological makers of organ damage were determined using the colorimetric kits indicated in the KRT according to the manufacturer’s instructions. All absorbance readings were performed in 96-well plates using an Infinite M200 plate reader (Tecan).

### Histopathology

Mouse liver, lung, and kidney were collected 30h after infection and immediately fixed in 10% buffered formalin. Samples were then embedded in paraffin, sectioned (3 μm) and stained for hematoxilin and eosin according to standard procedures. Blind histopathology analysis was performed by a trained pathologist at the Instituto Gulbenkian de Ciência Histopathology Unit. Tissues were scored for damage, namely necrosis and leukocyte infiltration.

### Transmission Electron microscopy

Mice were euthanized 24h after doxycycline treatment, perfused with 10 mL cold PBS through the left ventricle, followed by perfusion with 10 mL 2% formaldehyde. The gastrocnemius muscle was excised, cut in small pieces and fixated for 1 hour in 2% formaldehyde and 2.5% glutaraldehyde in 0.1 M phosphate buffer pH 7.4. Secondary fixation was performed with 1% osmium tetroxide for 30 min, followed by staining with 1% tannic acid for 20 min and 0.5% uranyl actetate for 1 hour. Samples were then dehydrated in a graded series of ethanol and embedded in Embed-812 epoxy resin. Sections (70 nm) were made using a Leica UC7 ultramicrotome and picked on slot grids coated with 1% formvar in chloroform. Samples were then post-stained with 1% uranyl acetate for 7 minutes and Reynolds lead citrate for 5 min. Transmission electron microscopy images were acquired on a Hitachi H-7650 microscope operating at 100 KeV and equipped with a XR41M mid mount AMT digital camera.

### Immunoblotting

Cultured cells were rinsed with PBS and lysed with RIPA buffer containing protease and phosphatase inhibitor cocktails. Frozen tissue samples were homogenized in RIPA buffer containing protease and phosphatase inhibitor cocktails using TissueLyser II. Cell and tissue homogenates were centrifuged at 20000 x*g* for 10 min at 4°C, the supernatant was collected and proteins quantified by the Bradford method. SDS-PAGE was performed by loading 20 μg total protein onto 10 or 12% polyacrylamide gel. Proteins were then transferred onto nitrocellulose membranes, blocked with 5% low-fat milk and incubated with primary antibodies for 16h at 4°C. HRP-conjugated secondary antibodies were incubated for 1h at room temperature and developed with ECL Prime. Chemiluminescence was acquired with GE Amersham Imager 680. Band density was analyzed with Fji version 1.52n.

### Gene expression analyses

#### RNA extraction and qPCR

For lung RNA-Seq, both lungs were harvested, cleaned from fat and bronchi and homogenized in 1 mL Trizol using a TissueLyser II. Liver samples (∼50 mg) were homogenized in 500 μL Trizol. Homogenates from both tissues were centrifuged at 20000 x*g* for 3 minutes at 4°C and 500 μL supernatant were used for RNA extraction. Extraction was performed with 100 μL chloroform and the aqueous layer was transferred to an RNeasy mini spin column. RNA purification was performed according to the manufacturer’s protocol including one step of in-column DNase treatment. RNA was quantified in Nanodrop and 1 μg total RNA was used to synthesize cDNA using SuperScript II and Oligo dT. Real-time quantitative PCR was performed using Sybr Green reagent and ABI QuantStudio 7 equipment.

#### RNA-Seq

Total RNA samples were checked for quality using AATI Fragment Analyzer. Only samples with RNA Quality Number (RQN) >7 and clearly defined 28S and 18S peaks were considered for downstream analysis.

mRNA libraries were prepared, pooled and sequenced (75 bp, single end) using NextSeq500.

#### Quality Assessment and Alignment

Prior to alignment, quality of the sequences was assessed using FASTQC and MultiQC (Ewels et al., 2016). Sequences were then aligned against the *Mus musculus* genome version 97, with the annotation file for the genome version 97, both obtained through the website of Ensembl. The alignment was performed using STAR (Dobin et al., 2013), with default parameters and with the option of *GeneCounts*.

#### Data analysis

The files obtained from *GeneCounts* were imported to R (version 3.5.3), taking into account the strandness inherent to the sequencing protocol. Downstream analysis was performed using DESeq2 (version 1.22.2) (Love et al., 2014). Data from raw counts were normalized through a Regularized Log Transformation (rlog) to create the Principal Component Analysis plot and Heatmaps (Love et al., 2014). The log2FC provided by the standard DESeq2 model was shrunk using the ‘ashr’ algorithm (Stephens, 2017). Gene Information was obtained using the package *org.Mm.eg.db*. For the purposes of this study, genes were considered differentially expressed when the p-value, adjusted using false discovery rate (FDR), was below 0.05.

### Liver metabolomics

#### Sample preparation

Liver samples (30-80 mg) were weighed and homogenized in 500 μl ice-cold methanol:acetonitrile:H2O (2:2:1, v/v) using a TissueLyser II. Homogenates were incubated at −80 °C for 4 hours and centrifuged at 20000 xg for 10 minutes at 4°C. The supernatant containing soluble fractions was stored at −80°C. The pellet was resuspended in 400 μl ice-cold 80% (v/v) methanol by vortexing for 1 minute at 4°C. Samples were then incubated for 30 minutes at −80°C and centrifuged at 20000 xg for 10 minutes at 4°C. Supernatant was collected and combined with the previously obtained supernatant containing soluble fractions. Samples were centrifuged again and the supernatant stored at −80°C until analysis.

#### Untargeted metabolomics

Extracted samples were thawed on ice, centrifuged for 2 min at 15,000 xg, and diluted according to the different sample weight with 0.1% formic acid (RP, reversed phase) or 50% acetonitrile (ACN) (HILIC, hydrophilic interaction chromatography). 2.5 μL of each diluted sample were pooled and used as a quality control (QC) sample. Samples were randomly assigned into the autosampler and metabolites were separated on a SeQuant ZIC-pHILIC HPLC column (Merck, 100 x 2.1 mm; 5 µm) or an RP-column (Waters, ACQUITY UPLC HSS T3 150 x 2.1; 1.8 μm) with a flow rate of 100 µl/min delivered through an Ultimate 3000 HPLC system (Thermo Fisher Scientific). The gradient ramp up time takes 25 min from 10% to 80% B in HILIC (A: ACN; B: 25 mM ammonium bicarbonate (ABC) in water) and from 1% to 90% B in RP (A: 0.1% FA in water; B: 0.1% FA in ACN). Metabolites were ionized via electrospray ionization in polarity switching mode after HILIC separation and in positive polarity mode after RP separation. Sample spectra were acquired by data-dependent high-resolution tandem mass spectrometry on a Q-Exactive Focus (Thermo Fisher Scientific). Ionization potential was set to +3.5/-3.0 kV, the sheet gas flow was set to 20, and an auxiliary gas flow of 5 was used. Samples were analyzed in a randomized fashion and QC samples were additionally measured in confirmation mode to obtain additional MS/MS spectra for identification. Obtained data sets were processed by compound discoverer 3.0 (Thermo Fisher Scientific). Compound annotation was conducted by searching the mzCloud database with a mass accuracy of 3 ppm for precursor masses and 10 ppm for fragment ion masses as well as ChemSpider with a mass accuracy of 3 ppm using BioCyc, Human Metabolome Database, KEGG, MassBank and MetaboLights as databases.

#### Targeted metabolomics

Each sample was injected onto a SeQuant ZIC-pHILIC HPLC column (Merck, 100 x 2.1 mm; 5 µm) operated with an Ultimate 3000 HPLC system (Dionex, Thermo Fisher Scientific) at a flow rate of 100 µl/min and directly coupled to a TSQ Quantiva mass spectrometer (Thermo Fisher Scientific). For all transitions, the optimal collision energy was defined by analyzing pure metabolite standards. Chromatograms were manually interpreted using TraceFinder (Thermo Fisher Scientific), validating experimental retention times with the respective quality controls of the pure substances. In HILIC a 15 minutes gradient (A: 95% ACN, 5% 10 mM aqueous ammonium acetate; B: 50% ACN 50% 10 mM aqueous ammonium acetate) was used for separation. The following transitions have been used for quantitation in the negative ion mode (2.8 kV): stearic acid 283.1 *m/z* → 265.1 *m/z* and palmitic acid 255.1 *m/z* → 227.1 and in the positive ion mode (3.2 kV): stearoylcarnitine 428.1 *m/z* → 85 m/z, palmitoylcarnitine 400.1 *m/z* → 85 *m/z*, myristoylcarnitine 372.1 *m/z* → 85 *m/z* and lauroylcarnitine 344.1 *m/z* → 85 *m/z*.

### Electron transport chain (ETC) complex activity

Enzymatic activity of ETC complexes in mouse liver was performed as previously described (Spinazzi et al., 2012). Briefly, frozen liver samples (50-100 mg) were homogenized in 1mL homogenization buffer containing 8 mM Tris, 16 mM KCl, 0.8 mM EGTA and 250 mM sucrose using TissueLyzer II. Lysates were centrifuged at 20000 x*g* for 10 min at 4°C, the supernatant collected and proteins quantified by the Bradford method. Samples were then diluted in homogenization buffer for a final concentration of 1 mg/mL. Complex I (NADH:ubiquinone oxidoreductase) activity was determined by following oxidation of 100 μM NADH at 340 nm in the presence of 50 mM KPi pH 7.5, 60 μM ubiquinone, 3 mg/mL BSA, 300 μM potassium cyanide (KCN), and 20 μg total protein. Complex II (succinate dehydrogenase) activity was determined by measuring the reduction of 80 μM dichloroindophenol sodium salt hydrate (DCPIP) at 600 nm in a reaction mixture containing 25 mM KPi pH 7.5, 20 mM succinate, 50 μM decylubiquinone, 1 mg/mL BSA, 300 μM KCN, and 10 μg total protein. Complex III (ubiquinol:cytochrome c oxidoreductase) activity was determined by measuring reduction of 75 μM cytochrome C at 550 nm in the presence of 25 mM KPi pH 7.5, 100 μM decylubiquinol (obtained by reduction of decylubiquinone with potassium borohydride), 0.025% (v/v) tween-20, 100 μM EDTA, 500 μM KCN, and 1.5 μg total protein. Complex IV (cytochrome c oxidase) activity was determined by measuring oxidation of 60 μM cytochrome C (previously reduced with sodium dithionite) in the presence of 50 mM KPi pH 7.0 and 1.0 μg total protein. Citrate synthase (CS) activity was determined by following reduction of 100 μM 5,5′-Dithiobis(2-nitrobenzoic acid) (DTNB) at 412 nm in a reaction mixture containing 100 mM Tris-HCl pH 8.0, 300 μM acetyl CoA, 0.1% (v/v) Triton X-100, 300 μM oxaloacetic acid, and 3 μg total protein. All reactions were performed at in 96-well plates (maximum of 12 simultaneous reactions) and absorbance was recorded using an Infinite M200 plate reader (Tecan). Unspecific activity of each complex was determined by performing a reaction in the presence of an appropriate inhibitor (rotenone for complex I, malonate for complex II, antimycin for complex III, and KCN for complex IV), which was then subtracted from the total activity of each sample. Enzymatic activity was calculated in nmol.min^.1^.mg protein^-1^, normalized for CS activity and expressed as percentage of the control.

### Flow cytometry

HepG2 cells were incubated with doxycycline 24h, tripsinyzed and centrifuged at 200 x*g* for 5 minutes at room temperature. Cells were then stained with 50 nM Mitotracker Green FM (for total mitochondrial content) for 30 minutes at 37°C, or with 5 μM MitoSOX Red (for mitochondrial ROS) for 10 minutes at 37°C in PBS supplemented with 2% FBS and 10 mM EDTA. Cells stained with MitoSOX were centrifuged, washed and resuspended in buffer, while cells stained with Mitotracker Green were centrifuged and resuspended without washing. Flow cytometry data were acquired on FACSCalibur (Becton Dickinson) and analyzed using the FlowJo software package (version 887).

## Quantification and statistical analysis

Mantel-Cox test was used for survival curve analysis. For infections with *E. coli*, mice with no changes in body temperature and weight within the first 24h (temperature >35°C and body weight>95%) were excluded from the analysis.

Mann-Whitney test was used for pairwise comparisons and two-way ANOVA with Tukey test was used for multiple comparisons. Statistical analysis was performed with Graphpad Prism 6.0 (GraphPad Software). The number of subjects used in each experiment is defined in figure legends. The following symbols were used in figures to indicate statistical significance: p <0.05 (*); p<0.01 (**); p<0.001 (***); p<0.0001 (****).

---

**Figure S1, related to Figure 1.**
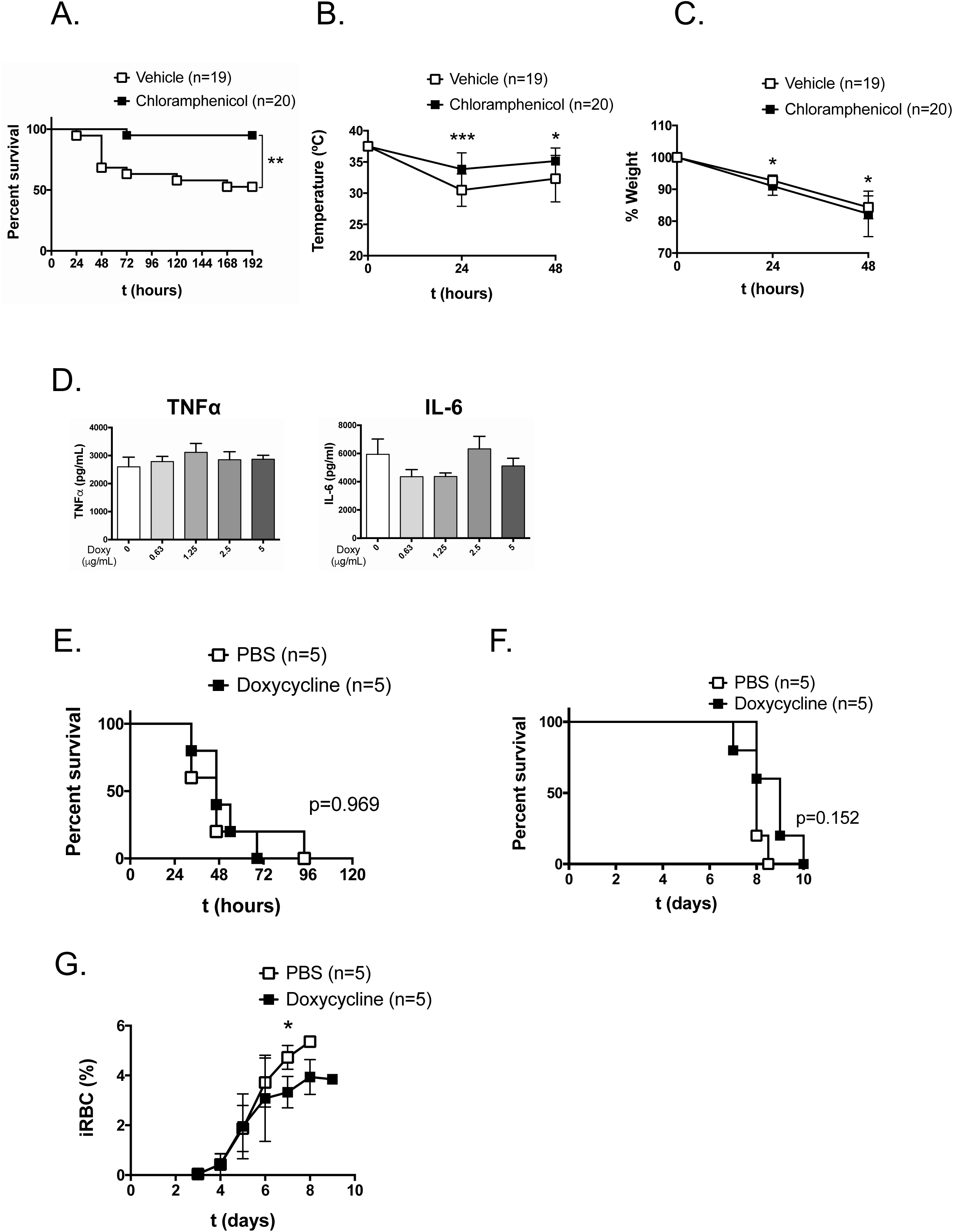
Characterization of doxycycline and chloramphenicol effects *in vivo* and *in vitro*. Survival (A), rectal temperature (B) and % initial body weight (C) after infection with TetR CamR *E. coli* and treatment with 50 μg/g body weight chloramphenicol (or 5% cyclodextrin as a control) at the time of infection. (D) TNFα and Interleukin (IL)-6 levels in supernatant of bone marrow-derived macrophages incubated with doxycycline for 1 h followed by stimulation with PFA-fixed *E. coli* (MOI=20) for 4 h. (E) Survival of mice infected with 5×10^5^CFU/mouse *Candida albicans* and treated with 1.75 μg/g body weight doxycycline (or PBS as a control) at 0, 24, and 48h. Survival (F) and parasitemia (G) levels (% infected red blood cells, *iRBC*) in mice infected with 1×10^5^/mouse *Plamodium berghei* Anka and treated with 1.75 μg/g body weight doxycycline (or PBS as a control) daily from days 0 to 8 post-infection. (A), (B), and (C) represent pooled data from three independent experiments. (D) represents a single experiment assayed in triplicate. (E), (F), and (G) represent a single experiment. Mean±SD are shown in (B), (C), (D), and (G).

**Figure S2, related to Figure 2.**
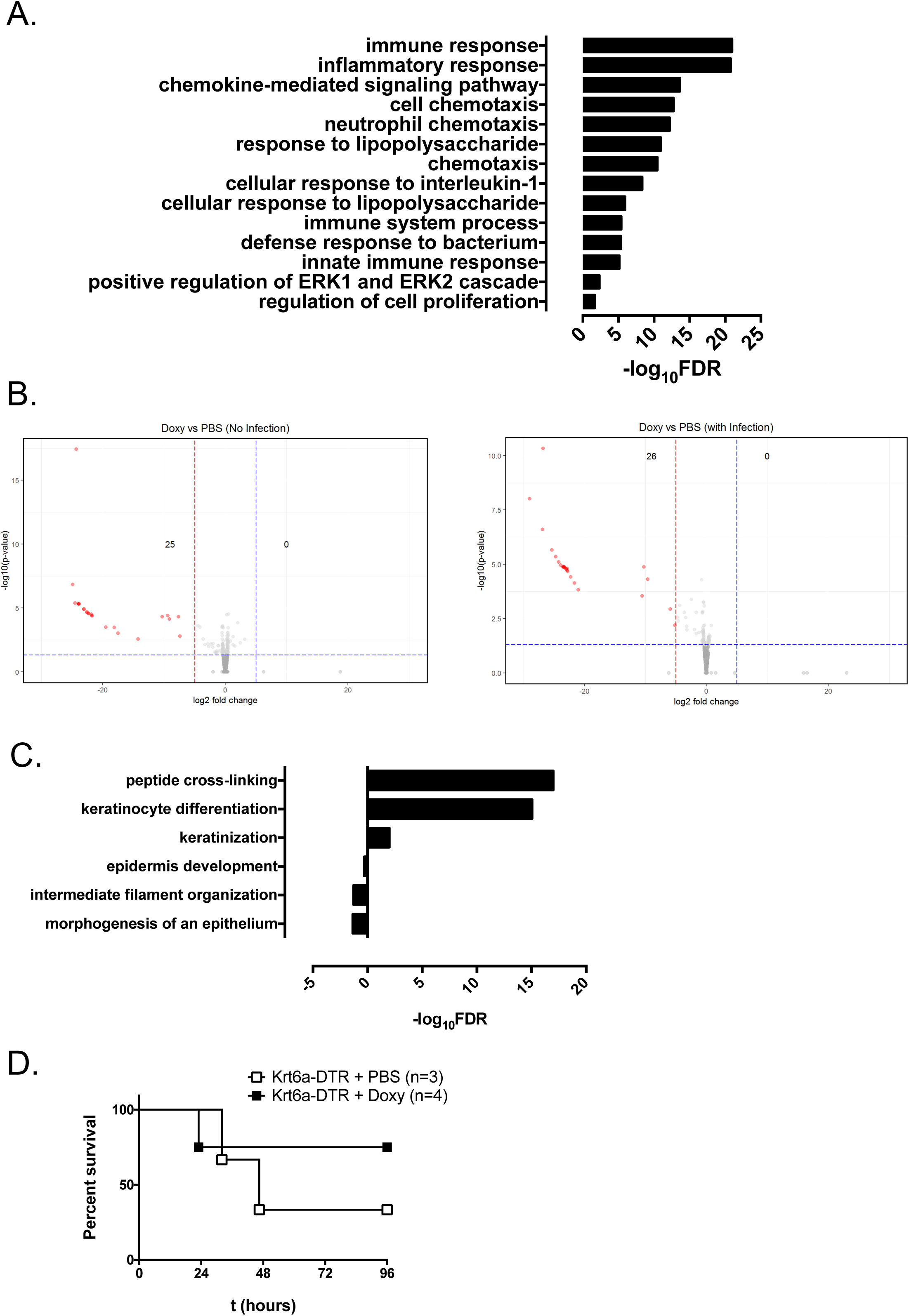
Lung RNA-Seq analysis. (A) Top GO_BP annotation of genes up-regulated during infection (PBS-treated mice), log2 fold change>5; p<0.05. (B) Volcano plots of doxycycline versus PBS-treated mice in the absence or presence of infection. Numbers indicate genes with log2 fold change <-5 or >5 and p<0.05. (C) Top GO_BP annotation of genes down-regulated with doxycycline treatment (non-infected mice), log2 fold change<-5; p<0.05. (D) Survival of Krt6a-depleted mice upon infection with *E. coli* and treatment with doxycycline. Krt6a-DTR mice were treated with 12 ng/g diphteria toxin by intra-tracheal administration 4 days before infection. Data represents a single experiment.

**Figure S3, related to Figure 3.**
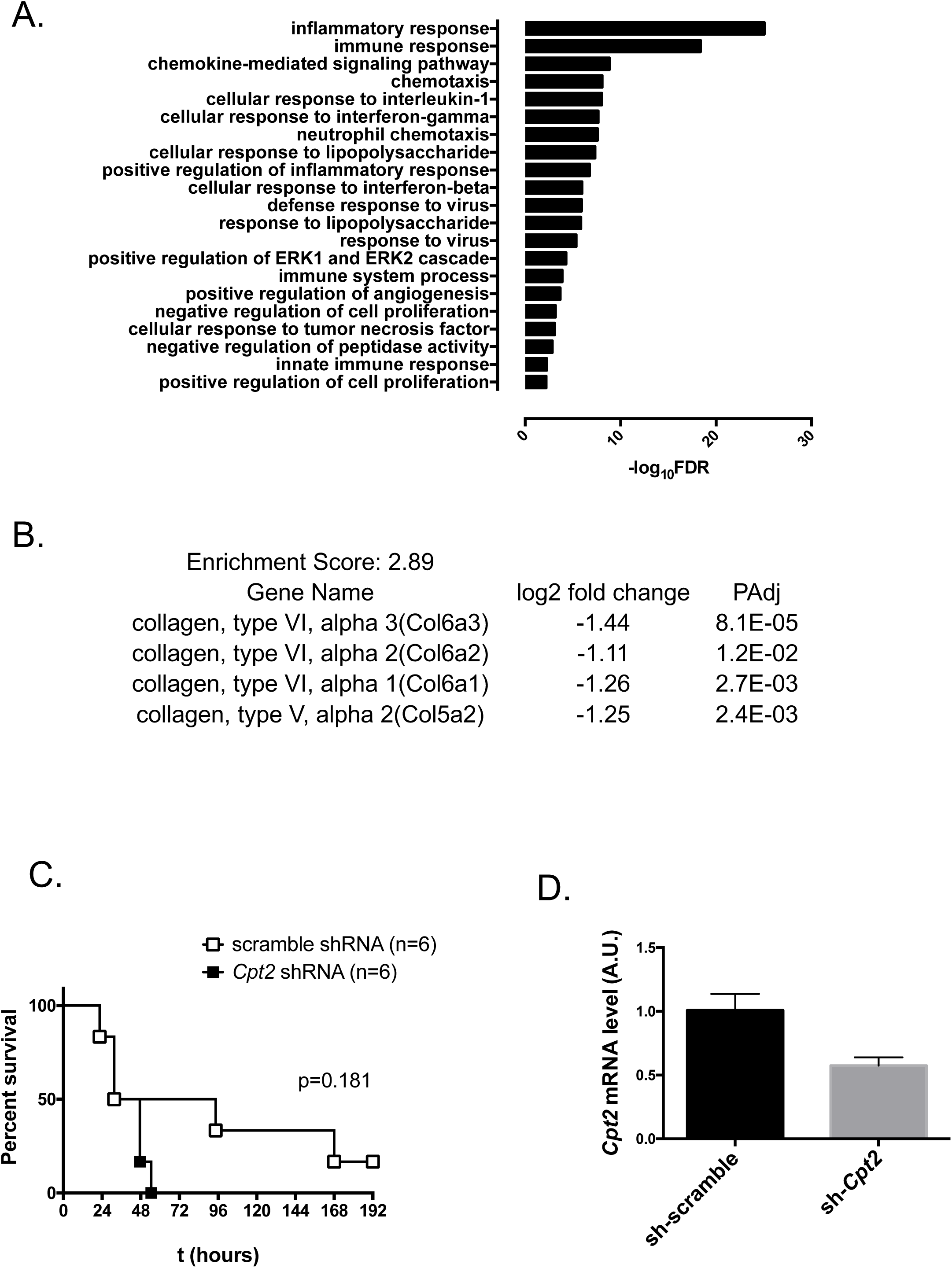
RNA-Seq analysis in liver. (A) Top GO_BP annotation of genes up-regulated during infection (PBS-treated mice), log2 fold change>5; p<0.05. (B) Gene functional clustering of down-regulated genes upon doxycycline treatment (infected groups), log2 fold change <-1, p<0.05. (C) Survival of C57BL/6J mice after infection with *E. coli*. Mice were injected with AAV8 expressing *Cpt2* shRNA (or scramble shRNA as a control) 7 days before infection. (D) *Cpt2* mRNA levels in mouse liver 7 days after injection of AAV8 *Cpt2* shRNA (or scramble shRNA as a control).

**Figure S4, related to Figure 4.**
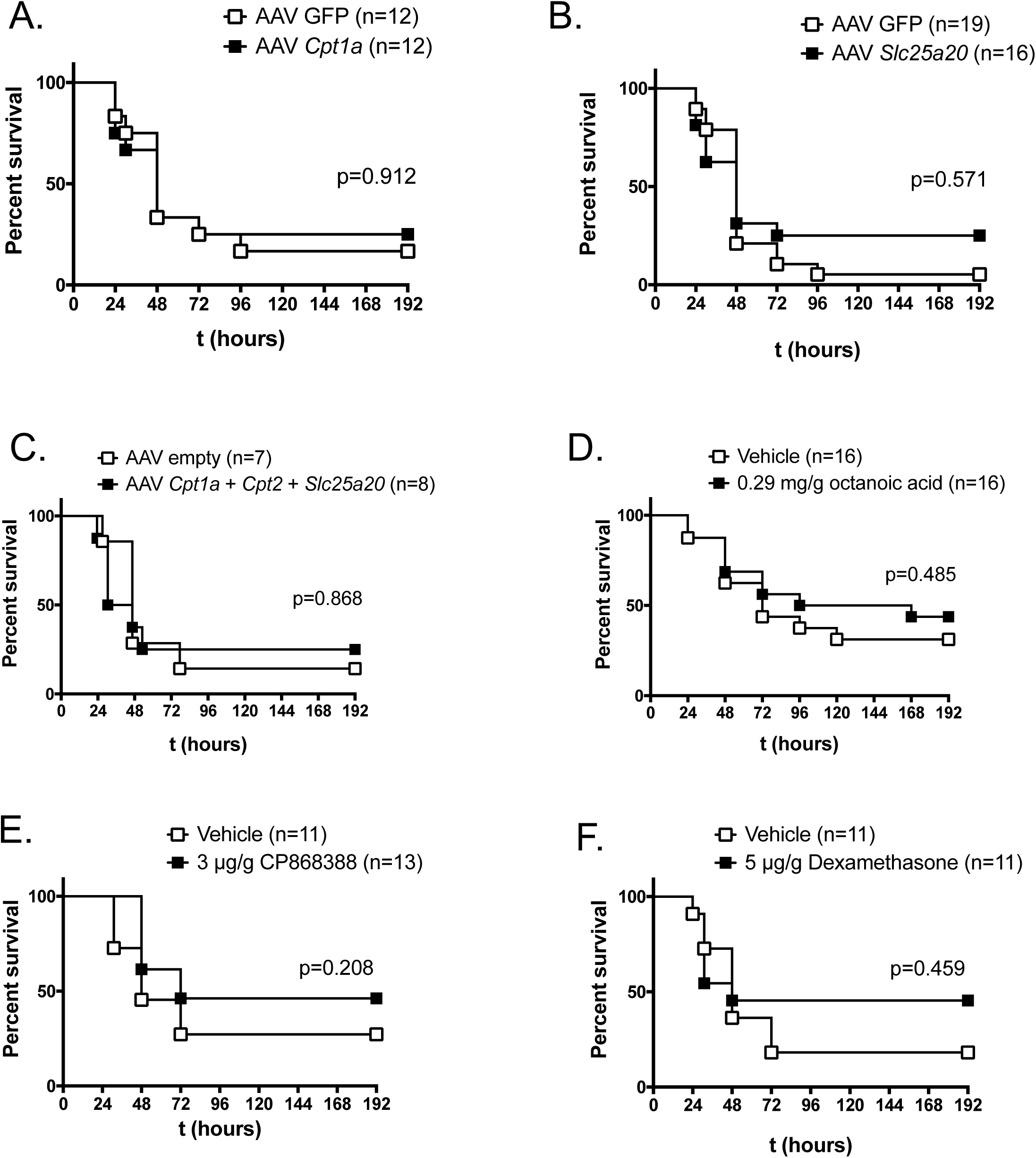
Fatty acid oxidation and glucocorticoid signaling gain-of-function experiments. (A, B, C) Survival of C57BL/6J mice after infection with *E. coli*. Mice were injected with AAV8-TBG expressing SLC25A20 (A), CPT1a (B) or a combination of CPT1a+CPT2+SLC25A20 (1:1:1) (C) one week before infection. (D) Survival of C57BL/6J mice after infection with *E. coli* and treatment with octanoic acid by oral gavage at 2, 8, 24, and 48h after infection. (E, F) Survival of C57BL/6J mice after infection with *E. coli* and treatment with CP868388 (PPARα agonist) (E) or dexamethasone (GR agonist) (F) by IP injection at the time of infection. (A) And (D) Represent pooled data from three independent experiments. (B), (E) and (F) represent pooled data from two independent experiments. (C) represents data from a single experiment.

**Figure S5, related to Figure 5.**
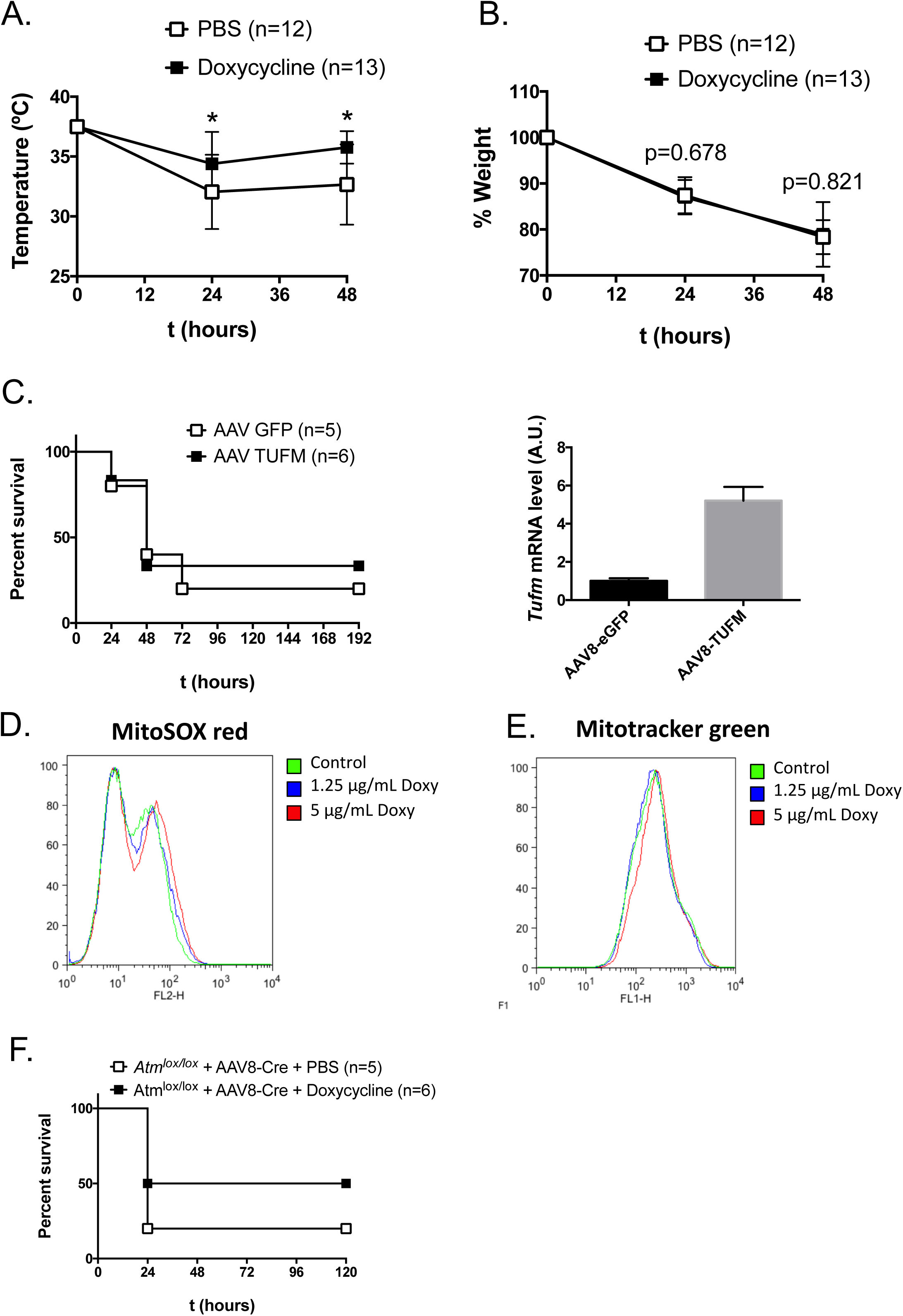
Possible mechanisms of doxycycline-induced disease tolerance. Rectal temperature (A) and % initial body weight (B) after infection of germ-free mice with TetR CamR *E. coli* and treatment with 1.75 μg/g body weight doxycycline (or PBS as a control) at 0, 24 and 48 h. Data represent mean±SD pooled from four independent experiments. (C) Survival of *E. coli*-infected mice and *Tufm* mRNA levels in mouse liver. Mice were injected with AAV8-TBG expressing TUFM one week before infection/liver collection. Data represents a single experiment. (D, E) Mitochondrial peroxide levels (MitoSOX red, D) and total mitochondrial content (Mitotracker green, E) in HepG2 cells after incubation for 24h with the indicated concentrations of doxycycline. Representative graphs of two independent experiments. (F) Survival of mice with liver-specific ATM knockdown upon infection with *E. coli* and treatment with doxycycline. *Atm*^lox/lox^ mice were treated with AAV8-Cre one week before infection. Data represents a single experiment.

**Figure S6, related to Figure 6.**
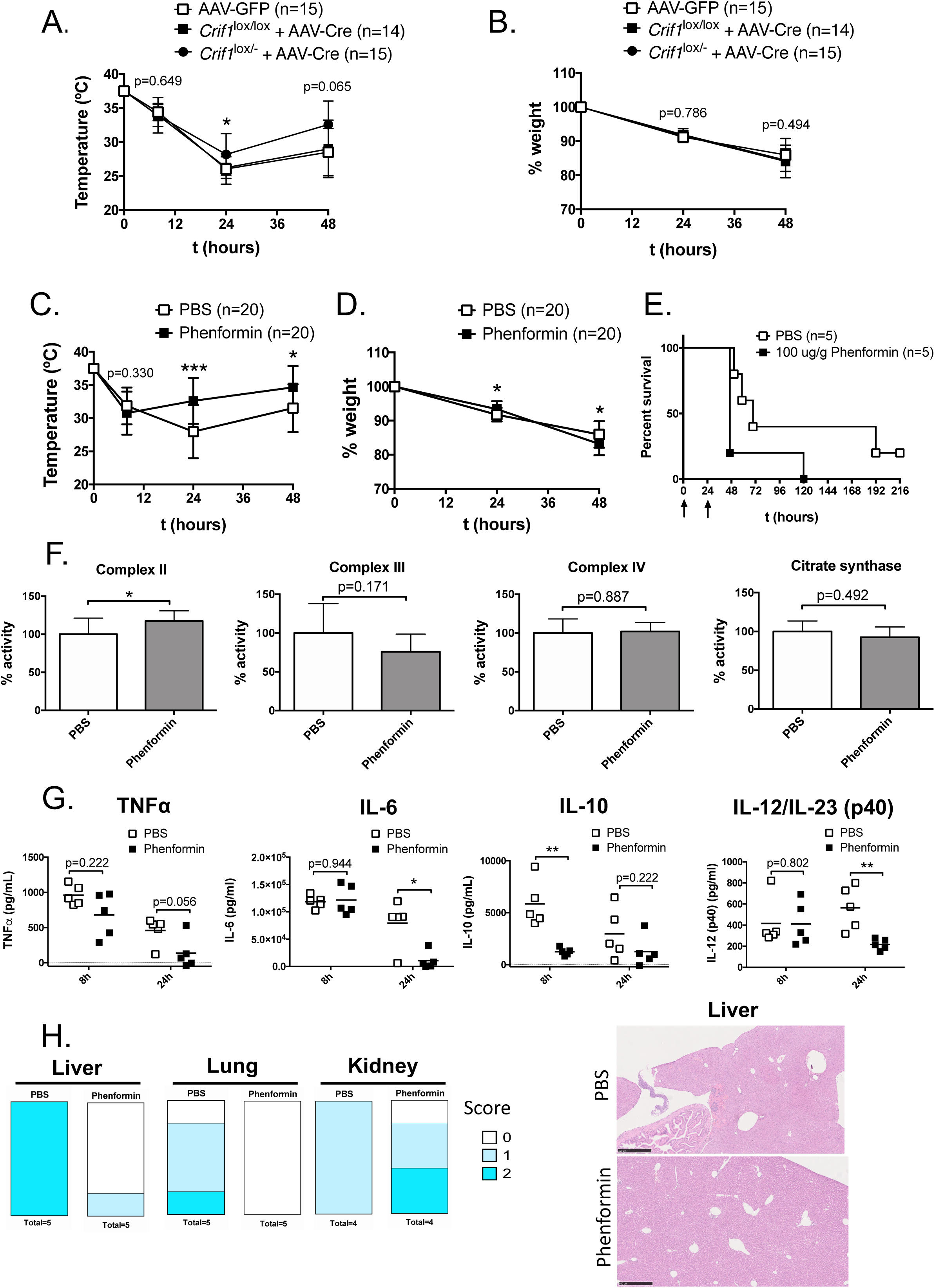
Protective effects of phenformin treatment and partial CRIF1 deletion. Rectal temperature (A) and % initial body weight (B) after infection of *Crif1*^lox/lox^ or *Crif1* ^lox/-^ mice with TetR CamR *E. coli*. Mice were previously injected with AAV8-TBG expressing Cre recombinase (or GFP as a control). Rectal temperature (C) and % initial body weight (D) after infection of C57BL/6J mice with *E. coli* and treatment with a single administration of 100 μg/g body weight phenformin (or PBS as a control) at the time of infection. (E) Survival after infection of C57BL/6J mice with *E. coli* and treatment with100 μg/g body weight phenformin (or PBS as a control) at 0 and 24h after infection. (F) Enzymatic activity of the ETC complexes II, III, IV and CS in mouse liver collected 12h after PBS or phenformin treatment. ETC activity is expressed as % of PBS-treated control, normalized for CS activity. (G) TNFα, IL-6, IL-10, and IL12/IL23(p40) levels in mouse serum at the indicated time-points after infection. (H) Organ damage score and representative images of Hematoxylin-Eosin stained slides from liver, lung, and kidney 24 h after infection. Score 0 = no lesions; 1 = very mild; 2 = mild lesions. (A) and (B) represent pooled data from three independent experiments. (C) and (D) represent pooled data from four independent experiments.(E)) Represents data from a single experiment. (F) Represents mean±SD of 5 mice/group from a single experiment. (G) Represents data from a single experiment; squares represent individual mice and bars indicate the mean.

